# Unique Features of Different Classes of G-Protein-Coupled Receptors Revealed from Sequence Coevolutionary and Structural Analysis

**DOI:** 10.1101/2021.06.03.446974

**Authors:** Hung N Do, Allan Haldane, Ronald M Levy, Yinglong Miao

## Abstract

G-protein-coupled receptors (GPCRs) are the largest family of human membrane proteins and represent the primary targets of about one third of currently marketed drugs. Despite the critical importance, experimental structures have been determined for only a limited portion of GPCRs and functional mechanisms of GPCRs remain poorly understood. Here, we have constructed novel sequence coevolutionary models of the A and B classes of GPCRs and compared them with residue contact frequency maps generated with available experimental structures. Significant portions of structural residue contacts were successfully detected in the sequence-based covariational models. “Exception” residue contacts predicted from sequence coevolutionary models but not available structures added missing links that were important for GPCR activation and allosteric modulation. Moreover, we identified distinct residue contacts involving different sets of functional motifs for GPCR activation, such as the Na^+^ pocket, CWxP, DRY, PIF and NPxxY motifs in the class A and the HETx and PxxG motifs in the class B. Finally, we systematically uncovered critical residue contacts tuned by allosteric modulation in the two classes of GPCRs, including those from the activation motifs and particularly the extracellular and intracellular loops in class A GPCRs. These findings provide a promising framework for rational design of ligands to regulate GPCR activation and allosteric modulation.

**Significance:** G-protein-coupled receptors (GPCRs) play key roles in cellular signaling and serve as the primary targets of ∼1/3 of currently marketed drugs. In this work, we have presented the first analysis of both residue sequence coevolution and structural contact maps in different classes of GPCRs. We have inferred pathways for GPCR signal transduction that could not be determined from structural analysis alone. Distinct residue contacts have been identified in the signaling pathways of class A and B GPCRs. Our combined sequence coevolutionary and structural contact analysis has thus revealed important insights into the mechanism of GPCR signal transduction, which is expected to facilitate rational drug design of the GPCRs.

## Introduction

G-protein-coupled receptors (GPCRs) comprise the largest and most diverse family of integral membrane proteins in eukaryotes. GPCRs mediate various physiological activities, including vision, olfaction, taste, neurotransmission, endocrine, and immune responses (1). Due to the critical roles in cellular signaling, approximately 34% of FDA-approved therapeutic agents act on GPCRs (2). On the basis of sequence homology and functional similarity, GPCRs are classified into six different classes, four of which are present in the human body: class A (Rhodopsin-like), class B (secretin receptors), which is further divided into subclasses of B1 (classical hormone receptors), B2 (adhesion GPCRs) and B3 (methuselah-type receptors), class C (metabotropic glutamate receptors) and class F (frizzled/TAS2 receptors) (3, 4). The other two classes include D (fungal mating pheromone receptors) and E (cyclic AMP receptors). In comparison, class A is the largest with 701 known receptors and by far the most extensively studied class of GPCRs (4). GPCRs share a characteristic structural fold of seven transmembrane (TM) α-helices (TM1-TM7) connected by three extracellular loops (ECL1-ECL3) and three intracellular loops (ICL1-ICL3). The extracellular and intracellular domains are typically important for binding of the ligands and G proteins, respectively. Consequently, the loop regions are notably diverse in sequences and structures.

Activation varies among the A and B classes of GPCRs. Class A GPCR activation is triggered by binding of an agonist to the receptor orthosteric pocket located within the 7TM domain (5). Upon agonist binding, the receptor intracellular end of TM6 moves outwards to open up an intracellular cavity to accommodate and activate the G protein (6–9). Activation of class B GPCRs requires binding of both the agonist and G protein, as well as disruption of the TM6 helix with a sharp kink (9–11).

Bioinformatics analysis has been previously carried out to identify important residue interactions for GPCR activation. Cvicek et al. generated a structure-based alignment of 25 GPCRs that was extended to include transmembrane sequences of all human GPCRs (12). The final sequence-structure alignment revealed 40 interhelical contacts that were common to class A GPCRs, 23 of which were conserved among class B, C, and F GPCRs. Furthermore, by comparing the active and inactive structures of class A receptors, they identified 15 Native ACtivation “Hot-spOt” residues (NACHOs) for class A GPCR activation (12). In 2019, Zhou et al. discovered a common activation pathway of class A GPCRs through an analysis residue-residue contact scores of 235 available class A GPCR structures (13). A four-layer activation pathway that connected the extracellular to intracellular regions was characterized at the residue level. Changes in critical residue contacts were identified during global movements of TM6 and TM7 in class A GPCR activation (13).

GPCRs are also able to bind allosteric ligands at topographically distinct sites, which could induce further conformational changes of the GPCRs (14). Allosteric ligands often include the positive allosteric modulator (PAM) and negative allosteric modulator (NAM) of GPCR activation. For class A GPCRs, binding of a PAM in the M2 muscarinic receptor was shown to induce slight contraction of the receptor extracellular pocket, which was pre-formed in the active agonist-bound structure (15, 16). Binding of a muscarinic toxin NAM to the inactive antagonist-bound M1 muscarinic receptor induced conformational changes in the receptor ECL2, TM1, TM2, TM6 and TM7 extracellular domains, as well as the TM2 and TM6 intracellular domains (17). In the free fatty acid receptor GPR40 and the C5a receptor, PAM binding in a lipid-facing pocket formed by TM3-TM4-ICL2 induced conformational changes in the ICL2, TM4 and TM5 of the active receptor (18, 19). The ICL2 adopted a short helical conformation and the TM5 was shifted along its helical axis towards the extracellular side relative to the TM4 (18). For class B GPCRs, a PAM was found to bind between the extracellular domains of TM1 and TM2 of the GLP-1 receptor (20). In the glucagon receptor, NAM binding restricted the outward movement of the TM6 intracellular domain (21). The ECL2 stretched to the central axis of the TM helical bundle, allowing for interactions from TM3 to TM6 and TM7 in the inactive class B GPCRs (21).

Despite remarkable advances in structural determination efforts, experimental structures have been resolved for only ∼90 unique GPCRs (3, 4). Functional mechanisms of many GPCR classes related to activation and allosteric modulation remain poorly understood at the residue level. Recent developments in methods for residue-covariation analysis have shown that another source of functional and structural information is in observed patterns of mutational covariation in multiple sequence alignments (MSAs) constructed from diverse protein sequences from a protein family (22–28). Residue covariation analysis methods infer a global probability model of sequences in the MSA which crucially captures the covariation of different columns of the MSA, while disentangling direct from indirect mutational covariation through inference of the underlying functional couplings which generated the observed covariation. The Direct Coupling Analysis or “Potts” models have been shown to capture important structural and functional information (24, 29–32). The models have further applications through the use of the probability model as a scoring function for individual sequences, for instance Potts models can predict effects of mutations to a sequence (33, 34), or be used to predict structural or conformational preferences of individual sequences or subfamilies of sequences (35, 36).

Here, we have constructed sequence coevolutionary Potts Hamiltonian models for class A and B GPCRs, for which sufficient protein sequences are available. We also generate residue contact frequency maps from available structures of both classes of GPCRs. Residue pairs that exhibit strong coevolutionary couplings but low structural contact frequencies are referred to as “exceptions” from the Potts model predictions. Several of such exception residue contacts added important missing links for activation and allosteric modulation of the GPCRs. We have also identified distinct residue contacts that are important for activation and allosteric modulation of these two classes of GPCRs.

## Results

### Structural residue contacts of GPCRs were detected in sequence coevolutionary models

We inferred separate sequence-based Potts coevolutionary models for the A and B classes of GPCRs and compared them with corresponding residue contact frequency maps (**Figure 1**). The input multiple sequence alignments (MSAs) included 84,481 sequences for class A and 17,804 sequences for class B GPCRs. Any sequences and columns in the MSAs with more than 10% gaps were removed, and through the correction to downweight phylogenetically similar sequences, we obtained 5126 and 902 effective sequences for class A and B GPCRs, respectively (**Table S1**). The residue sequences of class A and B GPCRs were also aligned using the *hmmalign* function of HMMER (37) (**Table S2**). Moreover, residue contact frequency maps were built from all available structures of each class of GPCRs (**Table S3**). The structures included the active, inactive, active agonist-PAM-bound and inactive antagonist-NAM-bound GPCRs for both classes as shown for a model class A GPCR in **Figure 1A** and a model class B GPCR in **Figure 1D**. The Potts coupling parameters, which were used to calculate Frobenius norms as residue contact scores, were visualized as a heatmap for each of the two classes of GPCRs. The Potts model was then directly compared with the residue contact frequency map of class A (**Figure 1B**) and class B (**Figure 1E**) of GPCRs.

**Figure 1.**
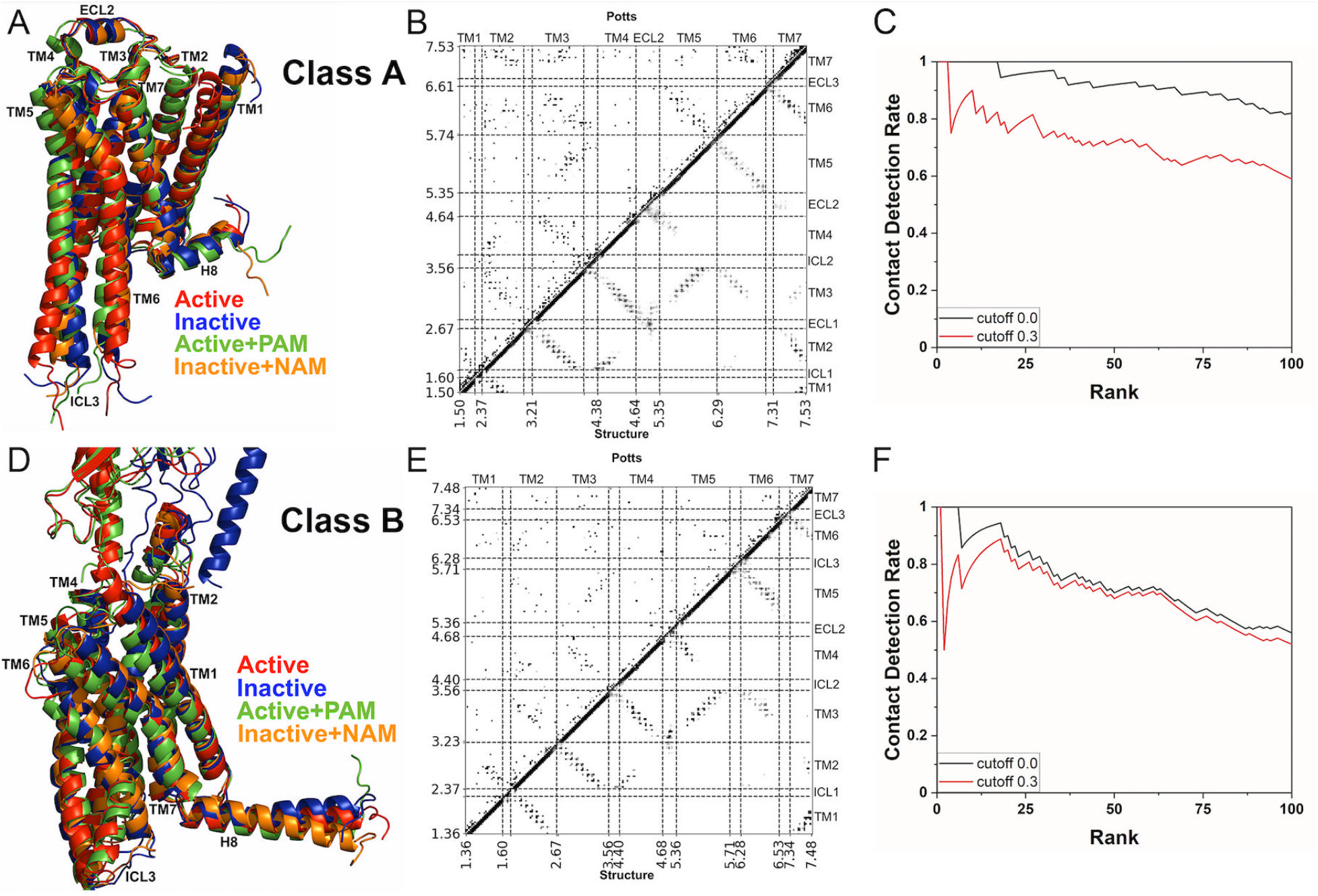
Comparison of the Potts model and residue contact frequency maps of class A and B GPCRs. **(A)** The active (red), inactive (blue), active agonist-PAM-bound (green) and inactive antagonist-NAM-bound (orange) structures of a model class A GPCR (PDBs: 6IBL, 5A8E, 6OIK and 6OBA). **(B)** Comparison of the Potts model (upper triangle) and residue contact frequency map (lower triangle) of class A GPCRs. **(C)** Structural contact detection rate of residue pairs in class A GPCRs with the top 100 long-distance Frobenius norms. **(D)** The active (red), inactive (blue), active agonist-PAM-bound (green) and inactive antagonist-NAM-bound (orange) structures of a model class B GPCR (PDBs: 6LMK, 5XEZ, 6VCB and 5EE7). **(E)** Comparison of the Potts model (upper triangle) and residue contact frequency map (lower triangle) of class B GPCRs. **(F)** Structural contact detection rate of residue pairs in class B GPCRs with the top 100 long-distance Frobenius norms. Only residue pairs with at least five residues apart were considered to calculate the contact detection rates using cutoffs of 0.0 and 0.3 for the contact frequency.

The correspondence between the Potts model and residue contact frequency map of class A GPCRs was evident in residue interactions of the TM6-TM7, TM5-TM6, TM3-TM6, TM3-TM5, TM3-TM4, TM2-TM7, TM2-TM4, TM2-TM3, TM1-TM7 and TM1-TM2 domains (**Figure 1B**). Overall, the number of predicted residue contacts from Potts model of class B GPCRs was lower than that of class A GPCRs. Nevertheless, the Potts model and residue contact frequency map of class B GPCRs were in agreement for residue interactions in the TM6-TM7, TM5-TM6, TM3-TM5, TM3-TM4, TM2-TM4, TM2-TM3, TM1-TM7 and TM1-TM2 domains (**Figure 1E**).

Furthermore, we extracted the top 100 long-distance Frobenius norms, consisting of residue pairs that were ≥ 5 residues apart in the numbering scheme, from the Potts model of each GPCR class and examined whether the corresponding residue contacts were observed in the experimental GPCR structures. The residue contacts predicted from the sequence-based Potts models were deemed true positives if they were present in available GPCR structures above a percentage cutoff (e.g., 30% or 0.3). Those residue pairs with significantly high Frobenius norms (particularly ≥0.16 for class A and ≥0.19 for class B) but with low contact frequencies below the percentage cutoff in the GPCR structures were considered exceptions in the Potts model predictions. A number of them were found to form important contacts for GPCR activation and allosteric modulation. With the cutoffs of 0.0 and 0.3 for the structural contact frequency, the residue contact detection rates of our Potts models were plotted for classes A and B of GPCRs in the **Figures 1C** and **1F**, respectively. With zero contact frequency cutoff, the residue contact detection rates were 0.82 and 0.56 for the top 100 long-distance Frobenius norms in the Potts models of the classes A and B of GPCRs, respectively. The detection rates decreased with increasing cutoff of the structural contact frequency. At the 0.3 contact frequency cutoff, the residue contact detection rates decreased to 0.59 and 0.52 for the top 100 long-distance Frobenius norms in the Potts models of the classes A and B of GPCRs, respectively. In comparison, the detection rate from Potts model of the class B GPCRs was slightly lower than that of class A GPCRs, due to the relatively smaller number of effective sequences (**Table S1**). This was consistent with previous studies that contact detection rate in the Potts model correlated with the number of effective sequences in the MSA (38, 39). Based on the above findings, the 0.3 cutoff of structural contact frequency was used for further analysis.

### Activation and Allosteric Modulation of Class A GPCRs

For class A GPCRs, contact exceptions with the top 20 long-distance Frobenius norms are summarized in **Table S4**. Notably, residue pair C3.44-V5.57 (rank 14) had the contact frequency increased to 0.38 in only the inactive class A GPCR structures, while residue pairs Y3.51-F5.56 (rank 28) and F5.47-L6.49 (rank 60) had the contact frequencies increased to 0.38 and 0.49 in only the active class A GPCR structures, respectively. Since these residue contacts showed significantly higher contact frequencies in one of the GPCR functional states (active or inactive), they were considered contacts that were important for GPCR activation or inactivation.

We built residue contact frequency maps for the active and inactive class A GPCRs and calculated their difference by subtracting residue contact frequencies in the inactive GPCR structures from those in the active structures. The residue contact frequency difference map between active and inactive class A GPCR structures is shown in **Figure 2A**. The exception residue contacts of rank 14, 28 and 60 were shown in **Figure 2B**. Furthermore, we highlighted the top 30 residue contacts with the largest differences of contact frequencies in the inactive and active GPCR structures (**Table S5A**). The seven switching residue contacts in the list included S3.39-F6.44, I3.46-L6.37 and R3.50-L6.34 that were present in only the inactive GPCR structures (**Figure 2E**) and I3.46-Y7.53, Y5.58-I6.40, Y5.62-L6.37 and A5.65-A6.33 that were present in only the active GPCR structures (**Figures 2F-2H**). Overall, the residue contacts between TM1-TM7 (**Figure 2C**) and TM3-TM6 (**Figure 2E**) were significantly weakened, while a number of contacts between TM2-TM7 (**Figure 2D**), TM3-TM7 (**Figure 2F**) and TM5-TM6 (**Figures 2G-2H**) were strengthened upon class A GPCR activation.

**Figure 2.**
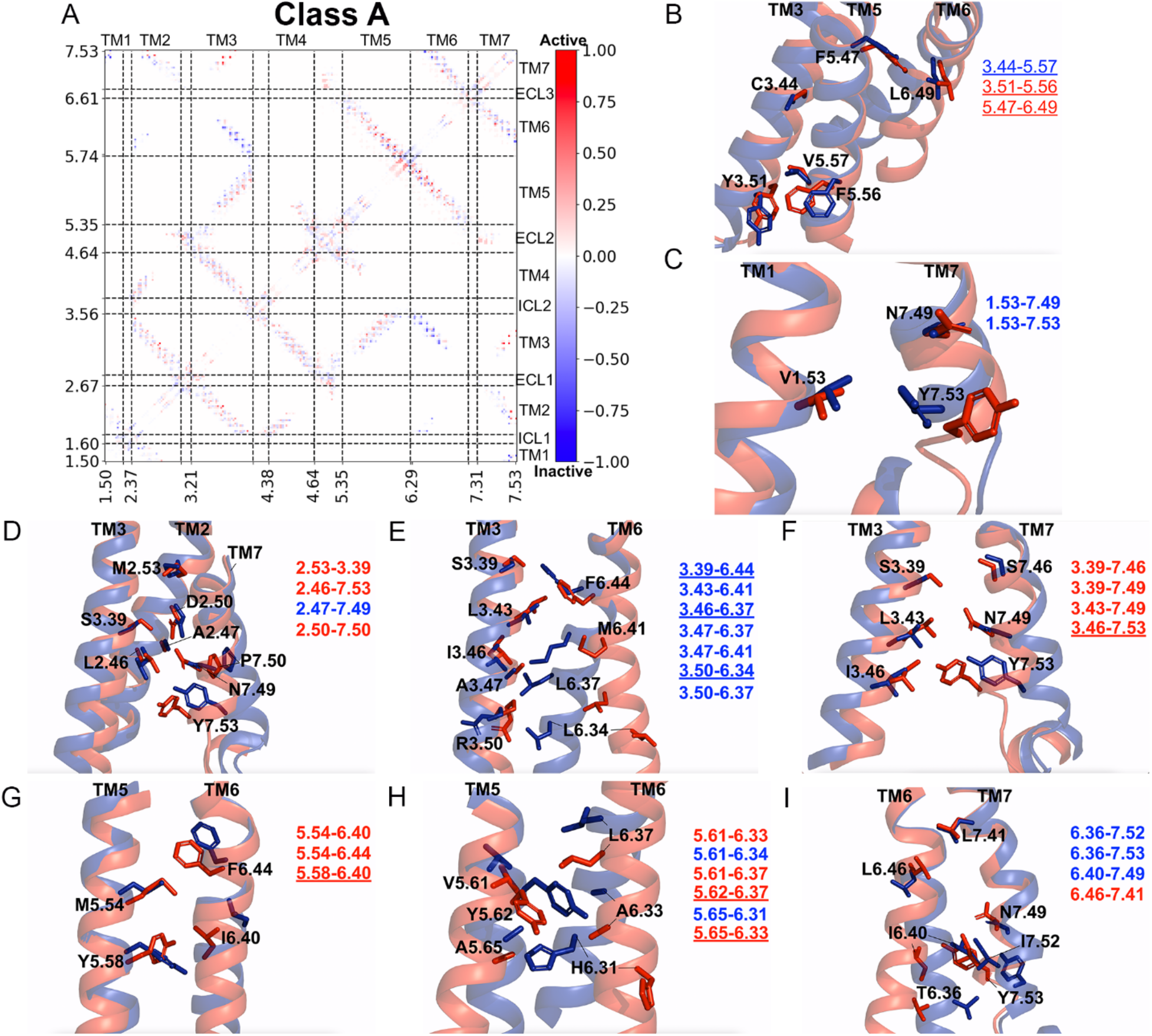
Residue contacts that are important for class A GPCR activation. **(A)** Comparison of residue contacts between the active and inactive structures of class A GPCRs. The residue contact frequency difference map was calculated by subtracting the residue contact frequencies in inactive structures from active structures. The residue contacts are colored by their differences in contact frequency in a blue (−1.00) – white (0.00) – red (1.00) color scale. **(B)** Exception residue contacts (rank 14, 28 and 60 in **Table S4**) between TM3-TM5 and TM5-TM6 that are changed during class A GPCR activation. **(C-I)** Residue contacts between TM1-TM7, TM2-TM7, TM3-TM6, TM3-TM7, TM5-TM6 and TM6-TM7 that showed the largest differences in contact frequencies between active and inactive class A GPCR structures. The exception, repacking and switching residue contacts are shown in underlined regular, bold and underlined bold fonts.

In addition to activation, we examined residue contacts that were tuned by allosteric modulation of class A GPCRs. Among the list of top ranked exception residue contacts (**Table S4**), the Y3.51-S3.56 residue pair (rank 4) had the contact frequency decreased by 0.33 upon binding of NAMs to the inactive GPCRs. With binding of PAMs to the active GPCRs, two residue pairs A2.49-W4.50 (rank 11) and D3.49-Y89^ICL2^ (rank 30) had the contact frequency decreased by 0.67 and 0.33, respectively, while the F5.47-L6.49 residue pair (rank 60) had the contact frequency increased by 0.33. These exception residue contacts in the Potts model thus resulted from allosteric modulation of class A GPCRs.

We generated additional residue contact frequency maps for the active agonist-bound class A GPCRs in the absence and presence of PAMs and calculated their difference of residue contact frequencies (**Figure 3A**), similarly for the inactive antagonist-bound class A GPCRs in the absence and presence of NAMs (**Figure 3B**). The residue contacts that were significantly tuned in class A GPCRs in the presence of allosteric modulators were listed in **Table S5B**. During binding of PAMs to active class A GPCRs, residues N2.39-R3.50 and N2.39-C3.53 formed new contacts, whereas the exception residue pair A2.49-W4.50 in the Potts model lost contact (**Figure 3C**). Moreover, residues L3.27-L4.62, V3.34-L4.56, D3.49-A4.42 and V5.51-W6.48 also formed new contacts in the active class A GPCRs (**Figures 3D-3E**). With binding of NAMs to the inactive class A GPCRs, residues T2.37-K4.39 formed new contacts between TM2-TM4 (**Figure 3F**), similarly for residue contacts I3.31-S4.57 between TM3-TM4 (**Figure 3F**), contacts A2.49-V3.36 between TM2-TM3 (**Figure 3G**) and contacts A5.39-H6.58 between TM5-TM6 (**Figure 3H**).

**Figure 3.**
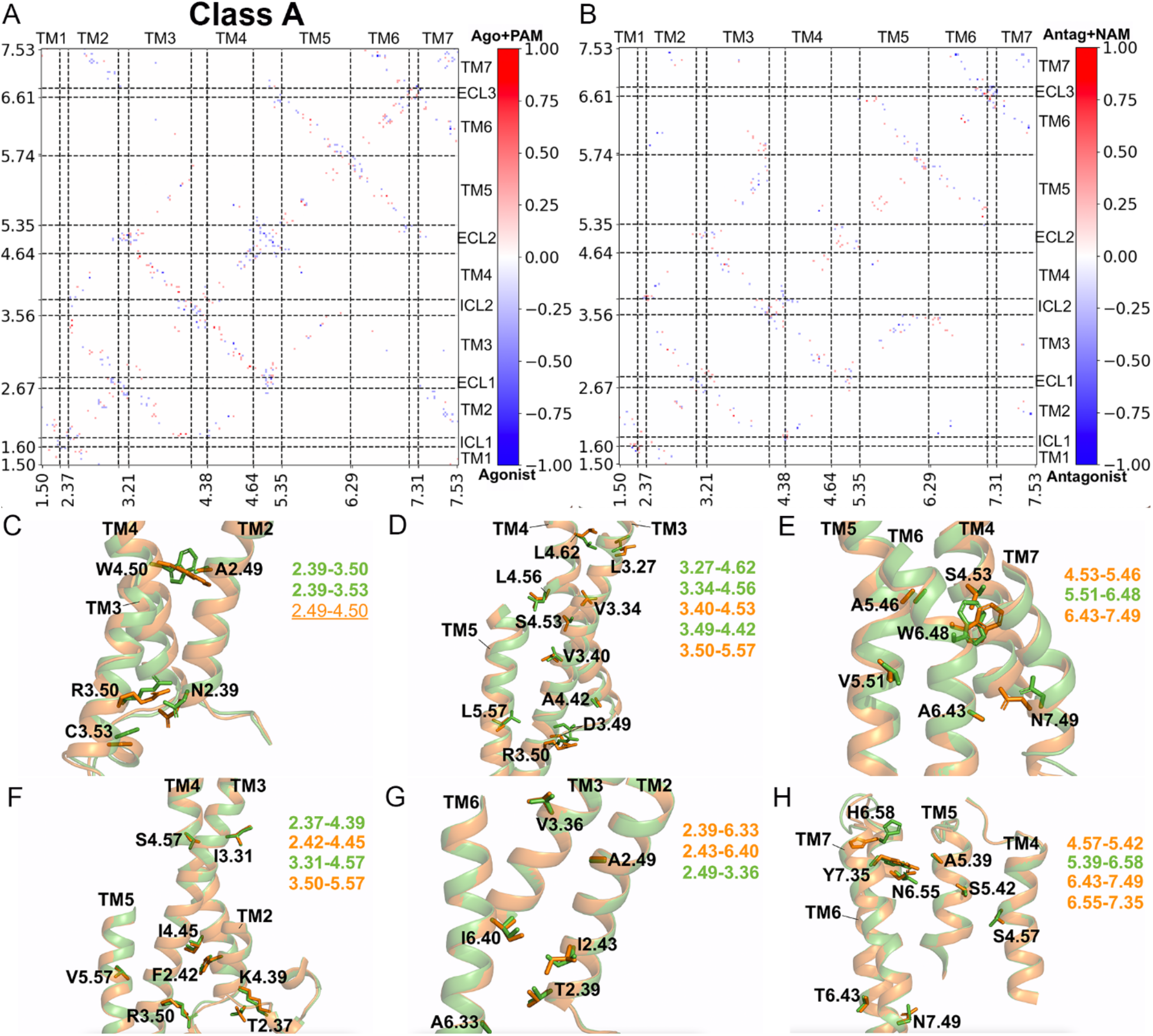
Residue contacts tuned by allosteric modulation in class A GPCRs. **(A-B)** Comparison of residue contacts between the agonist-PAM vs. agonist bound structures and antagonist-NAM vs antagonist bound structures of class A GPCRs. The residue contact frequency difference maps were calculated by subtracting the residue contact frequencies in structures without from structures with modulators. **(C-E)** Residue contacts between TM2-TM3, TM2-TM4, TM3-TM4, TM3-TM5, TM4-TM5, TM5-TM6 and TM6-TM7 that change upon binding of PAMs to the active agonist-bound class A GPCR structures. **(F-H)** Residue contacts between TM2-TM3, TM2-TM4, TM2-TM6, TM3-TM4, TM3-TM5, TM4-TM5, TM5-TM6 and TM6-TM7 that change upon binding of NAMs to the inactive antagonist-bound class A GPCR structure. Residue contacts that are strengthened and weakened are highlighted in green and orange, respectively. The exception residue contact is shown in underlined regular font.

### Activation and Allosteric Modulation of Class B GPCRs

For class B GPCRs, exception residue contacts with the top 20 long-distance Frobenius norms are summarized in **Table S6**. The contact frequency of residue pair S2.49-W4.50 (rank 2) increased to 0.31 in the active class B GPCR structures, while only 0.14 in the inactive structures. Therefore, this exception residue contact predicted in the Potts model could be explained by activation of class B GPCRs.

We computed the residue contact frequency difference map between active and inactive class B GPCR structures as shown in **Figure 4A**. The rank 2 exception residue contact is shown in **Figure 4B**. We highlighted the top 30 residue contacts with the largest differences in contact frequencies between active and inactive class B GPCR structures (**Table S7A**). A sharp kink was formed in the middle of the TM6 helix, leading to switching contacts including L3.43-L6.43, E3.46-L6.43, E3.46-L6.44, G3.47-L6.44, N5.54-L6.44, F5.58-L6.44, V5.62-T6.37/L6.40, L5.65-6.33 and L5.69-6.33 in the active class B GPCR structures (**Figures 4C, 4F** and **4G**). Meanwhile, the TM6 intracellular end moved away from the TM3, TM5 and TM7 helices during class B GPCR activation, losing a number of key residue contacts that were formed in the inactive receptor structures, including L3.50-L6.33/A6.34/T6.37, L3.54-K6.30, L3.43-6.40/I6.41, E3.46-T6.37/L6.40, I5.61-A6.34, L6.40-L7.48 and L6.44-Q7.45 (**Figures 4D, 4E** and **4H**).

**Figure 4.**
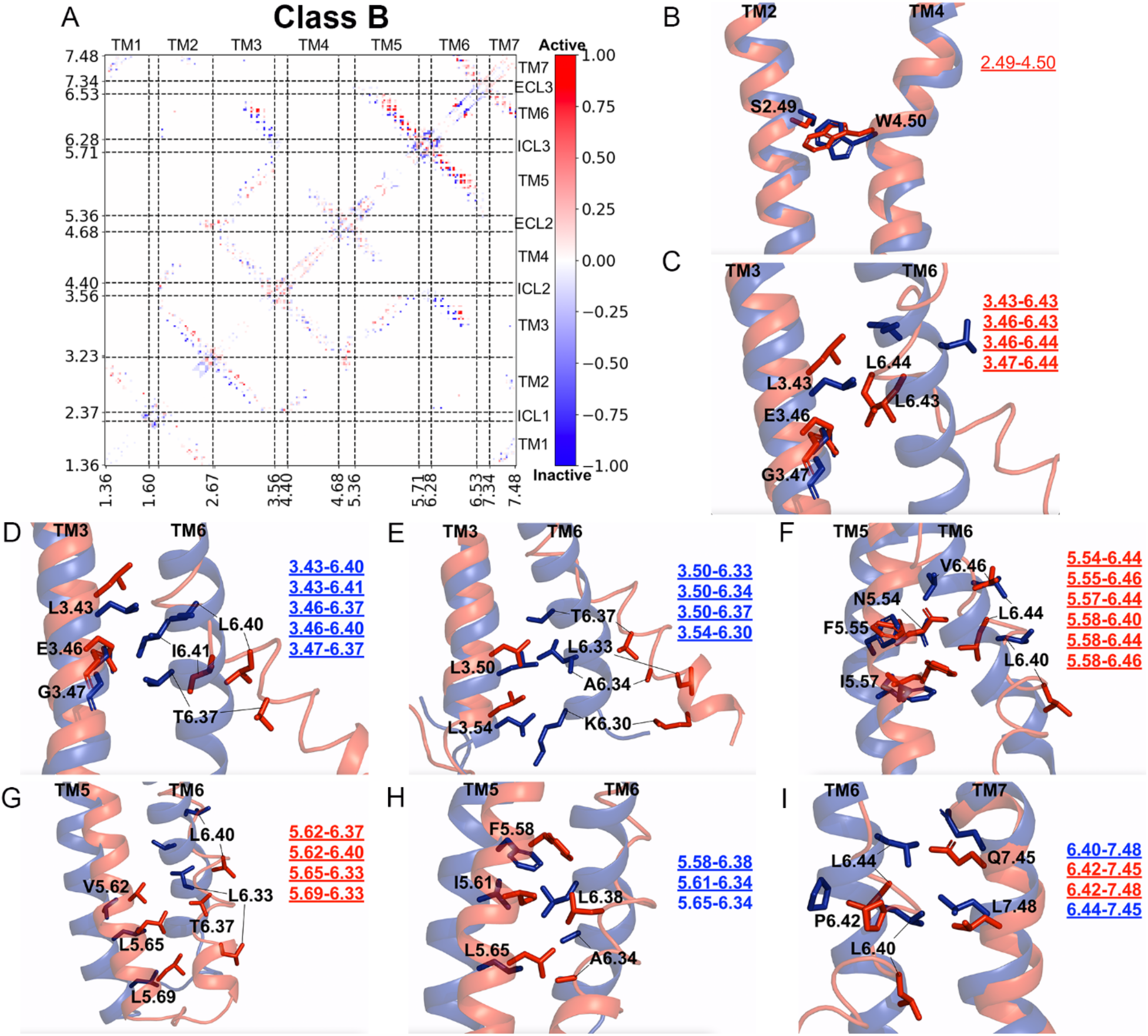
Residue contacts that are important for class B GPCR activation. **(A)** Comparison of residue contacts between the active and inactive structures of class B GPCRs. The residue contact frequency difference map was calculated by subtracting the residue contact frequencies in inactive structures from active structures. The residue contacts are colored by their differences in contact frequency in a blue (−1.00) – white (0.00) – red (1.00) color scale. **(B)** The exception residue contact (rank 2 in **Table S6**) between TM2-TM4 that are changed during class B GPCR activation. **(C-I)** Residue contacts between TM3-TM6, TM5-TM6 and TM6-TM7 that showed the largest differences in contact frequencies between active and inactive class B GPCR structures. The exception, repacking and switching residue contacts are shown in underlined regular, bold and underlined bold fonts.

We also computed the residue contact frequency difference map for the active agonist-bound class B GPCRs in the absence and presence of PAMs (**Figure 5A**), similarly for the inactive antagonist-bound class B GPCRs in the absence and presence of NAMs (**Figure 5B**). The residue contacts that were significantly tuned in the presence of allosteric modulators were listed in **Table S7B**. With the binding of PAMs in active class B GPCR structures, residue pairs G3.47-L6.43, L3.50-L6.43 and L3.50-L6.44 formed new contacts between TM3-TM6 (**Figure 5C**), similarly for residue contacts I5.51-V6.46, L5.52-A6.52, N5.54-V6.46 and P5.55-L6.44 between TM5-TM6 (**Figure 5D**) and residue contacts H6.47-T7.37, H6.47-T7.42, E6.48-E7.38 and I6.50-E7.38 between TM6-TM7 (**Figure 5F**). With the binding of NAMs in the inactive class B GPCR structures, residue pairs L3.43-T6.37, E3.46-I6.41 and L3.50-K6.30 formed new contacts between TM3-TM6 (**Figure 5G**), similarly for contacts W5.40-A6.52, R5.44-E6.48 and F5.48-A6.52 between TM5-TM6 (**Figure 5H**) and contacts H6.47-Q7.45, V6.50-K7.34 and V6.50-D7.38 between TM6-TM7 (**Figure 5I**).

**Figure 5.**
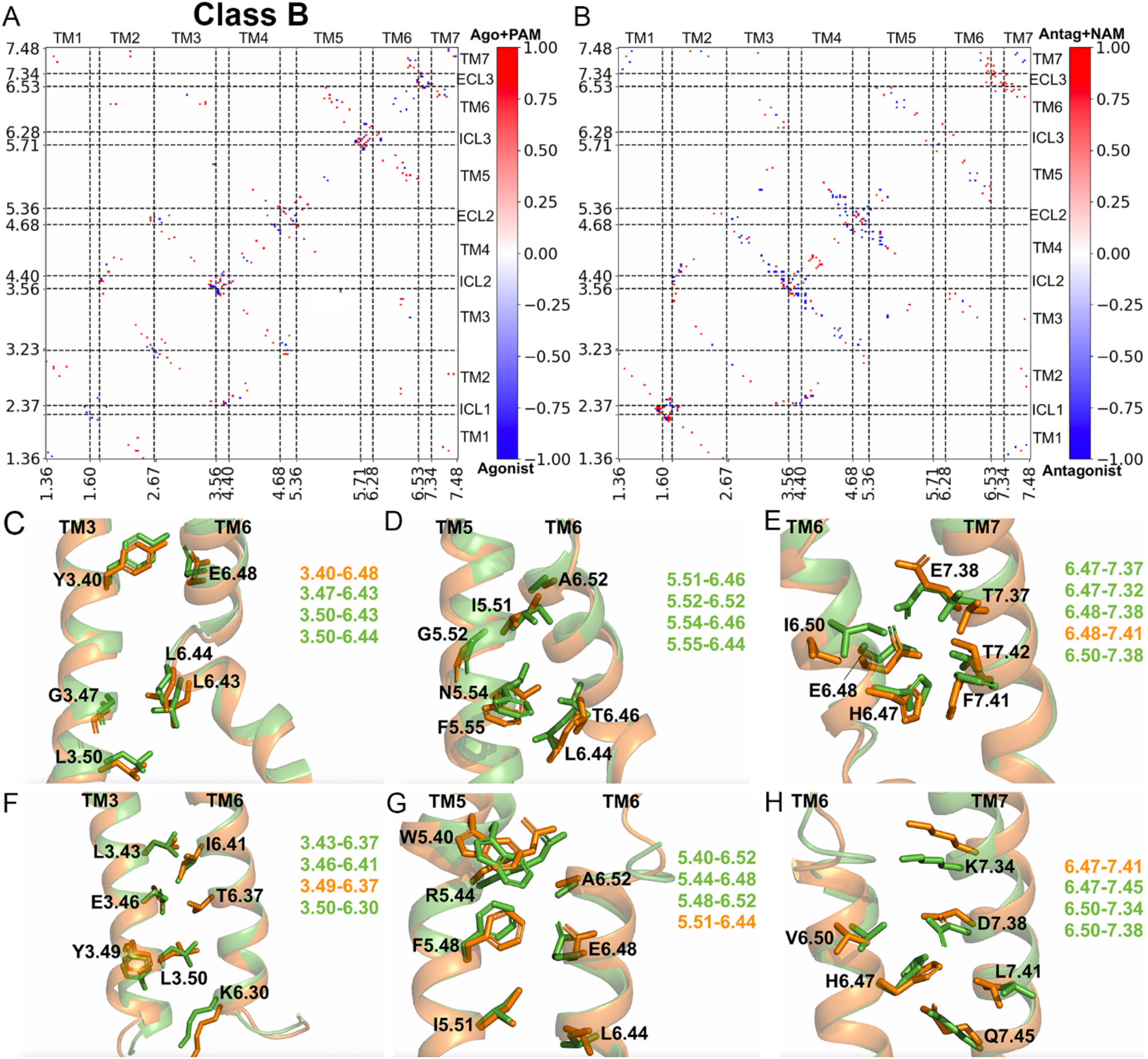
Residue contacts tuned by allosteric modulation in class B GPCRs. **(A-B)** Comparison of residue contacts between the agonist-PAM vs agonist bound structures and antagonist-NAM and antagonist bound structures of class B GPCRs. The residue contact frequency difference maps were calculated by subtracting the residue contact frequencies in structures without from structures with modulators. **(C-E)** Residue contacts between TM3-TM6, TM5-TM6 and TM6-TM7 that change upon binding of PAMs to the active agonist-bound class B GPCR structures. **(F-H)** Residue contacts between TM3-TM6, TM5-TM6 and TM6-TM7 that change upon binding of NAMs to the inactive antagonist-bound class B GPCR structures. Residue contacts that are strengthened and weakened are highlighted in green and orange, respectively.

### “Exception” residue contacts added important missing links for GPCR signaling

A number of exception contacts predicted from the Potts models added missing links that were critical for the activation and allosteric modulation of GPCRs (**Figure 6**) (33, 34). Specifically, 6 and 1 of the top 20 exception contacts resulted from activation and/or allosteric modulation of the class A and B GPCRs, respectively (**Tables S4** and **S6**). In class A GPCRs, exception contacts 3.51-5.56 and 5.47-6.49 augmented the residue network for receptor activation of by involving the DRY and CWxP functional motifs (40), respectively (**Figure 6A**). Exception contacts 3.49-149^ICL2^ and 3.51-3.56, both of which involved the DRY motif, were tuned off upon binding of PAMs and NAMs to the class A GPCRs, respectively (**Figures 6B** and **6C**). Another exception contact between 2.49 (Na^+^ pocket) and 4.50 was important for both allosteric modulation of class A GPCRs (41–44) and activation of class B GPCRs (**Figures 6B** and **6D**). This was consistent with previous findings that W4.50 is fully conserved in class A and B GPCRs and its mutation significantly affected activation of the class B GPCRs (1, 45). In addition, certain exception residue contacts in the Potts models appeared to result from arbitrary contact definition. They showed average Cβ-Cβ distances close to the 8Å contact cutoff, including 3 residue pairs T3.42-W4.50, D3.32-Y7.43 and L2.46-S7.46 in class A GPCRs and 1 residue pair I3.36-S7.42 in class B GPCRs (**Table S8**).

**Figure 6.**
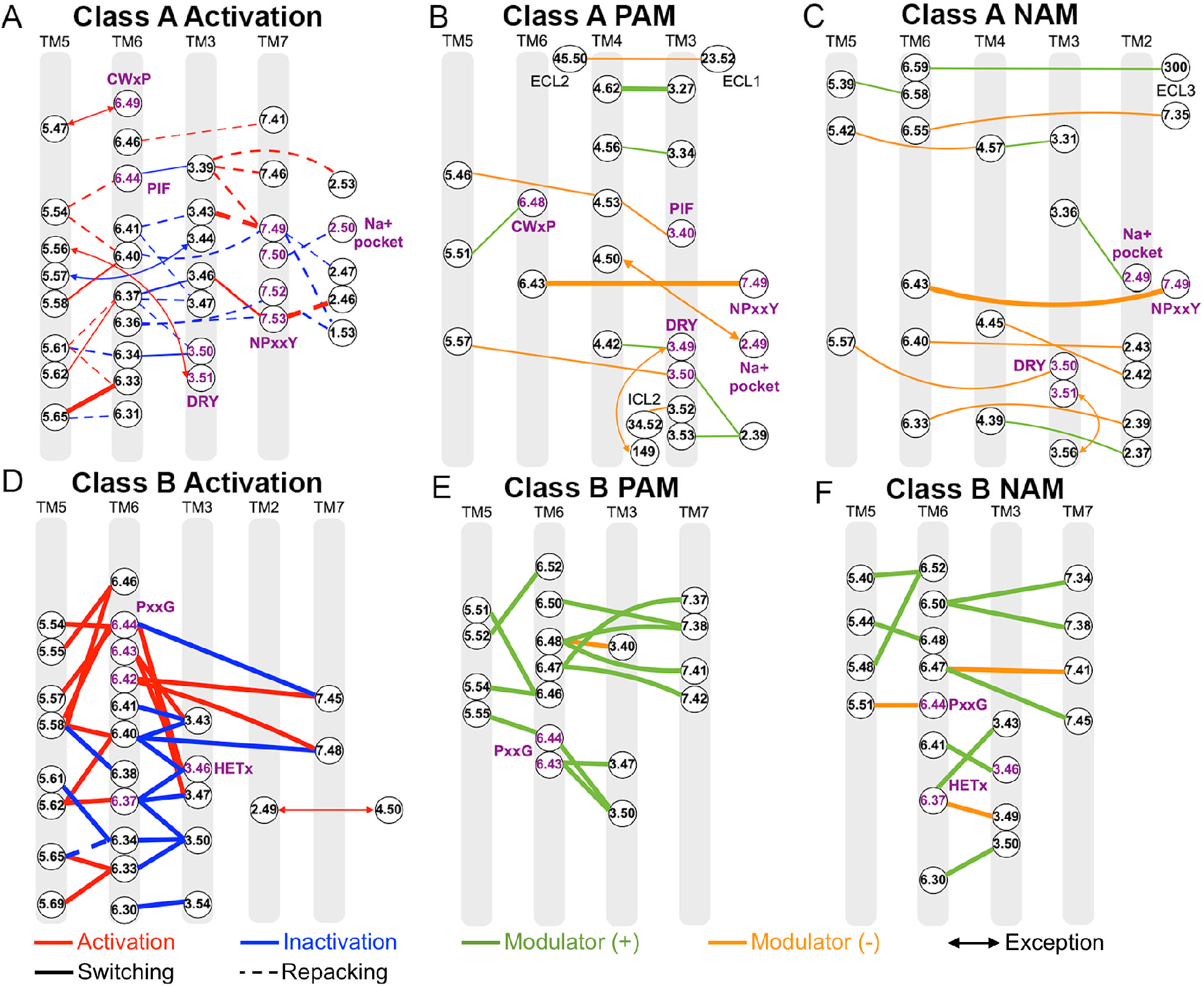
Summary of residue contacts tuned by activation and allosteric modulation in class A and B GPCRs. (**A-C**) Residue contacts that are tuned by activation and binding of PAMs and NAMs in class A GPCRs. (**D-F**) Residue contacts that are tuned by activation and binding of PAMs and NAMs in class B GPCRs. The conserved CWxP, PIF, DRY and NPxxY functional motifs in class A and the PxxG and HETx motifs in class B GPCRs are labeled. Contacts that are important for activation and inactivation are colored red and blue. Contacts that are formed in the presence and absence of allosteric modulators are colored green and orange, respectively. The line thickness is proportional to the magnitude of differences in residue contact frequencies (**Tables S5** and **S7**).

### Comparison of activation pathways of class A and B GPCRs

We have calculated residue contact frequency difference maps between active versus inactive structures to identify important residue contacts for GPCR activation. These maps systematically uncovered residue contacts that are important for activation of class A and B GPCRs. For the most extensively studied class A GPCRs, our results were mostly consistent with previous findings. Among our 30 residue pairs that showed the largest differences in contact frequencies between the active and inactive class A GPCR structures, 10 of them involved the previously identified NACHOs (12) and 9 of them were found in the common activation pathway of class A GPCRs (13) (**Table S9**). Five of the previous switching residue contacts (13) became repacking contacts in this analysis provided a significant increase in the number of available experimental structures. Our analysis also revealed 20 additional residue contacts involved in class A GPCR activation (**Table S9**). Upon activation of class A GPCRs, residue contacts were lost between the intracellular domains of TM6 and TM3 (involving the highly conserved DRY motif), as well as for the conserved NPxxY motif in TM7 with the TM1, TM2 (involving the Na^+^ binding pocket) (44, 46) and TM6. New residue contacts were formed between TM5-TM6 (involving the extracellular CWxP and PIF motifs) and TM3-TM7 (involving the NPxxY motif) (46) (**Figure 6A**). It is well established that the TM6 intracellular end moves outwards, while the highly conserved NPxxY motif in the TM7 intracellular domain moves towards TM3 (5, 46–52), which resulted in the switching TM3 residue contacts from TM6 to TM7. Our structural analysis determined the residue contacts that were weakened between TM1-TM7 and TM3-TM6 and strengthened between TM3-TM7 in this global movement.

For class B GPCR activation, all 30 top-ranked residue contacts involved residues from TM6, which is known to distort significantly to the point of losing helical properties and form a sharp kink in the middle with the intracellular domain pointing outwards (9, 11, 53). Due to dramatic conformational change of TM6, new switching contacts were formed near its sharp kink for the conserved PxxG motif, while residue contacts were lost for the TM3 (the HETx motif) and the TM6 intracellular domain (residues K6.30-I6.41) with TM5 and TM7 (**Figure 6D**).

### Comparison of allosteric modulation of class A and B GPCRs

Our sequence coevolutionary and structural contact analysis further revealed important residue contacts tuned by allosteric modulation of the A and B classes of GPCRs (**Figure 6**). Overall, the TM helices that were found important for activation could be also allosterically modulated in GPCRs. For class A GPCRs, residue contacts between TM2-TM3, TM2-TM4, TM3-TM4, TM3-TM5, TM4-TM5, TM5-TM6 and TM6-TM7 changed significantly upon binding of allosteric modulators (**Figures 6B** and **6C**). Residue pairs including 3.27-4.62, 3.34-4.56 and 5.51-6.48 (toggle switch in the CWxP motif) formed new contacts upon binding of PAMs, being consistent with the previous finding that PAM binding induced slight contraction of the receptor extracellular mouth (15, 54). Furthermore, residue contacts such as 4.53-5.46 and 5.51-6.48 were modified, due to shift of the TM5 towards the extracellular side relative to the TM4 (18). PAM binding also introduced rearrangements of residue contacts in the three extracellular loops and ICL2 of the class A GPCRs (**Figure 6B**). On the other hand, NAM binding induced conformational changes in the TM2 (the Na^+^ binding pocket) (41–44), TM3 (the DRY motif), TM4, TM5, TM6 and TM7 (the NPxxY motif) domains of the class A GPCRs (**Figure 6C**) (17, 19). For class B GPCRs, residues in the TM6, which is heavily involved in activation of the receptors (9, 21), were also found to undergo substantial changes of contacts in the presence of allosteric modulators, especially near the PxxG and HETx motifs (**Figures 6E** and **6F**). Our study, for the first time, systematically identified residue contacts that are tuned by allosteric modulation in different classes of GPCRs. They provide a framework for rationally designing selective allosteric drugs to modulate the structure and function of GPCRs.

## Discussion

We have revealed unique features of classes A and B GPCRs from combined sequence coevolutionary and structural contact analysis. A total number of 9 and 2 of the top 20 exception residue contacts in the coevolutionary models have been explained for class A and B GPCRs, respectively. They result from activation and allosteric modulation of GPCRs, as well as the arbitrary contact definition. However, the remaining exception residue contacts are still unexplained (**Tables S4** and **S6**). A certain number of these exceptions could possibly result from inaccuracy of the sequence coevolutionary models, especially for the class B GPCRs that has relatively smaller numbers of effective sequences. Further explanations of the other exception residue contacts, however, can be potentially obtained with more GPCR structures that are yet to be determined in the future (4).

The number of inactive experimental structures of class A GPCRs is about twice of their active structures (**Table S1**). More of the latter are thus needed, which has been recently boosted by remarkable advances in the cryo-EM technique for solving the active GPCR-G protein complex structures. More experimental structures are also needed for the class B GPCRs, especially the class B2 GPCRs for which the first structure has been obtained only very recently (55). Only a handful of structures are currently available for the GPCRs bound by allosteric modulators. Because the allosteric modulators are advantageous over traditional agonists and antagonists for providing more selective therapeutic drugs of GPCRs, a significantly larger number of allosteric modulator-bound structures may be expected in the near future(14). In addition, more structures associated with new functional mechanisms of GPCRs (e.g., biased agonism (56)) await to be determined. When more of the above structures become available, they should be periodically added for updated structural contact analysis. Different residue contacts can be potentially identified and used to explain the remaining exception contacts in Potts models of the GPCRs.

Compared with the inactive and active structures, intermediate structures have been rarely determined for GPCRs (**Table S1**). This largely results from the highly dynamic nature and relatively short lifetime of the GPCR intermediates, which have proven difficult to characterize in experiments. In this context, computational molecular dynamics simulations that have captured activation pathway and intermediate conformations of the GPCRs (7, 57–59) provide a promising approach to address the challenge. The simulation derived intermediate structures can be also used for additional structural contact analysis. Meanwhile, it will enable a more complete sequence-structure-dynamics-function relationship analysis of the GPCRs.

Furthermore, while the number of sequences for class C, D, E and F GPCRs are too small to construct accurate sequence coevolutionary models (38, 39), we can still perform structural contact analysis and dynamics simulations of these GPCRs. These studies will help us to identify important residue contacts and understand functional mechanisms of these even less studied or orphan GPCRs. Finally, it is worthy constructing a sequence coevolutionary model for the GPCR superfamily. When enough representative structures are obtained for each class of GPCRs, we can include all the structures for more complete analysis. Then the residue contact frequency map can be compared with the sequence coevolutionary model uncover common features of the different GPCR classes in the entire superfamily.

This analysis demonstrates how the combination of residue covariation and structural statistical analysis can supplement each other when applied to GPCR datasets. It establishes the basic connections between the sequence covariations and GPCR structural contacts, setting the stage for more advanced uses of the coevolutionary model to study GPCR function. Notably, the coevolutionary model has additional uses in predicting the energetics of individual sequences and particular residue pairs. For instance, the coevolutionary coupling values for residue pairs in a single target sequence have been used in “threading” calculations with protein structures to determine the propensity of that sequence to take on particular functionally active or inactive conformations (35). The coevolutionary model can also be used to predict functional characteristics of individual residue-residue interactions (60). Applying this sequence-based energetics view to the GPCR family may further uncover details of GPCR function. These analyses and detailed understanding of the functional mechanisms are expected to greatly facilitate rational structure-based drug design of the pharmaceutically important GPCRs.

## Methods

Multiple sequence alignments (MSAs) of class A and B GPCRs were built from Pfam IDs of PF00001 and PF00002, respectively. Sequences and columns in the MSAs with more than 10% gaps and phylogenetically related sequences at a similarity cutoff of 40% were removed. The Mi3-GPU (61) was applied to infer Potts coevolutionary models of class A and B GPCRs. Three rounds of inference were performed for each Potts model. In the first round, 2^18^ walkers with 64 MCMC steps were used to reduce the sum-of-square-residuals, average bivariate marginal relative error and covariance energy. The second round was performed using an increased number of 2^20^ walkers with 32-64 MCMC steps to fully level off the covariance energy values. The final round with 2^22^ walkers for four MCMC steps was performed to minimize the finite-sampling error and obtain a model with statistically accurate marginals and negligible residuals (61). In addition, residue contact frequencies were calculated from 283 structures of class A GPCRs (94 active, 185 inactive, and four intermediate) and 31 class B GPCR structures (16 active, 14 inactive, and one intermediate). Residue contact frequency difference maps were also built for active versus inactive GPCR structures to identify residue contacts involved in GPCR activation, as well as allosteric modulator-bound and modulator-free GPCR structures to identify residue contacts tuned by allosteric modulation in the class A and B GPCRs. Details of the MSA, sequence coevolutionary models and structural contact analysis are provided in **SI Methods**.

## Acknowledgements

This work used supercomputing resources with allocation awards TG-MCB180049 through the Extreme Science and Engineering Discovery Environment (XSEDE), which is supported by National Science Foundation (NSF) grant ACI-1548562 and project M2874 through the National Energy Research Scientific Computing Center (NERSC), which is a U.S. Department of Energy Office of Science User Facility operated under Contract No. DE-AC02-05CH11231, and computational resources provided by the Research Computing Cluster at the University of Kansas. This work was supported in part by the American Heart Association (Award 17SDG33370094 to Y.M.), the National Institutes of Health (R01GM132572 to Y.M. and 5U54-AI150472 and R35GM132090 to R.L.), NSF (193484 to R.L.) and the startup funding in the College of Liberal Arts and Sciences at the University of Kansas (to Y.M.).

## Author Contributions

A.H., R.M.L. and Y.M. designed the study. H.D. performed all calculations. H.D., A.H., R.M.L. and Y.M. analyzed data and wrote the manuscript.

## References

1. A. J. Venkatakrishnan et al., Molecular signatures of G-protein-coupled receptors. Nature 494, 185–194 (2013).

2. A. S. Hauser et al., Pharmacogenomics of GPCR drug targets. Cell 172, 41–54 (2018).

3. R. C. Stevens et al., The GPCR Network: a large-scale collaboration to determine human GPCR structure and function. Nature Reviews Drug Discovery 12, 25–34 (2013).

4. V. Isberg et al., GPCRdb: an information system for G protein-coupled receptors. Nucleic Acids Research 44, D365–364 (2016).

5. L. Ye, N. Van Eps, M. Zimmer, O. P. Ernst, R. S. Prosser, Activation of the A2A adenosine G-protein-coupled receptor by conformational selection. Nature 533, 265–268 (2016).

6. R. Nygaard et al., The dynamic process of beta(2)-adrenergic receptor activation. Cell 152, 532–542 (2013).

7. R. O. Dror et al., Activation mechanism of the β2-adrenergic receptor. Proc. Natl. Acad. Sci. U.S.A. 108, 18684–18689 (2011).

8. Y. Miao, S. E. Nichols, P. M. Gasper, V. T. Metzger, J. A. McCammon, Activation and dynamic network of the M2 muscarinic receptor. Proc Natl Acad Sci 110, 10982–10987 (2013).

9. D. Hilger et al., Structural insights into difference in G protein activation by family A and family B GPCRs. Science 369, eaba3373 (2020).

10. Y. Zhang et al., Cryo-EM structure of the activated GLP-1 receptor in complex with a G protein. Nature 546, 248 (2017).

11. Y. L. Liang et al., Toward a Structural Understanding of Class B GPCR Peptide Binding and Activation. Mol Cell 77, 656–668 e655 (2020).

12. V. Cvicek, W. I. Goddard, R. Abrol, Structure-Based Sequence Alignment of the Transmembrane Domains of All Human GPCRs: Phylogenetic, Structural, and Functional Implications. PLoS Computational Biology 12, e1004805 (2016).

13. Q. Zhou et al., Common activation mechanism of class A GPCRs. Elife 19, e50279 (2019).

14. D. M. Thal, A. Glukhova, P. M. Sexton, A. Christopoulos, Structural insights into G-protein-coupled receptor allostery. Nature 559, 45–53 (2018).

15. A. C. Kruse et al., Activation and allosteric modulation of a muscarinic acetylcholine receptor. Nature 504, 101–106 (2013).

16. S. Maeda, Q. Qu, M. J. Robertson, G. Skiniotis, B. K. Kobilka, Structures of the M1 and M2 muscarinic acetylcholine receptor/G-protein complexes. Science 364, 552–557 (2019).

17. S. Maeda et al., Structure and selectivity engineering of the M1 muscarinic receptor toxin complex. Science 369, 161–167 (2020).

18. J. Lu et al., Structural basis for the cooperative allosteric activation of the free fatty acid receptor GPR40. Nature Structural and Molecular Biology 24, 570–577 (2017).

19. H. Liu et al., Orthosteric and allosteric action of the C5a receptor antagonists. Nat Struct Mol Biol 25, 472–481 (2018).

20. A. B. Bueno et al., Structural insights into probe-dependent positive allosterism of the GLP-1 receptor. Nat Chem Biol 16, 1105–1110 (2020).

21. A. Jazayeri et al., Extra-helical binding site of a glucagon receptor antagonist. Nature 533, 274–277 (2016).

22. G. M. Suel, S. W. Lockless, M. A. Wall, R. Ranganathan, Evolutionarily conserved networks of residues mediate allosteric communication in proteins. Nat Struct Biol 10, 59–69 (2003).

23. R. I. Dima, D. Thirumalai, Determination of network of residues that regulate allostery in protein families using sequence analysis. Protein Science 15, 258–268 (2006).

24. F. Morcos et al., Direct-coupling analysis of residue coevolution captures native contacts across many protein families. Proceedings of the National Academy of Sciences 108, E1293 (2011).

25. S. E. Nichols, C. X. Hernandez, Y. Wang, J. A. McCammon, Structure-based network analysis of an evolved G protein-coupled receptor homodimer interface. Protein science : a publication of the Protein Society 22, 745–754 (2013).

26. R. M. Levy, A. Haldane, W. F. Flynn, Potts Hamiltonian models of protein co-variation, free energy landscapes, and evolutionary fitness. Current Opinion in Structural Biology 43, 55–62 (2017).

27. I. Anishchenko, S. Ovchinnikov, H. Kamisetty, D. Baker, Origin of coevolution between residues distant in protein 3D structures. Proc Natl Acad Sci U S A 114, 9122 (2017).

28. J. M. Nicoludis, R. Gaudet, Applications of sequence coevolution in membrane protein biochemistry. Biochim Biophys Acta Biomembr 1860, 895–908 (2018).

29. J. I. Sulkowska, F. Morcos, M. Weigt, T. Hwa, J. Onuchic, Genomics-aided structure prediction. Proc Natl Acad Sci U S A 109, 10340–10345 (2012).

30. D. S. Marks, T. A. Hopf, C. Sander, Protein structure prediction from sequence variation. Nature Biotechnology 30, 1072–1080 (2012).

31. F. Morcos, N. P. Schafer, R. R. Cheng, J. Onuchic, P. G. Wolynes, Coevolutionary information, protein folding landscapes, and the thermodynamics of natural selection. Proc Natl Acad Sci U S A 111, 12408–12413 (2014).

32. S. Ovchinnikov et al., Large-scale determination of previously unsolved protein structures using evolutionary information. Elife 4, e09248 (2015).

33. M. Figliuzzi, H. Jacquier, A. Schug, O. Tenaillon, M. Weigt, Coevolutionary Landscape Inference and the Context-Dependence of Mutations in Beta-Lactamase TEM-1. Molecular biology and evolution 33 (2016).

34. T. A. Hopf et al., Mutation effects predicted from sequence co-variation. Nature Biotechnology 35, 128–135 (2017).

35. A. Haldane, W. F. Flynn, P. He, R. S. Vijayan, R. M. Levy, Structural propensities of kinase family proteins from a Potts model of residue co-variation. Protein science : a publication of the Protein Society 25, 1378–1384 (2016).

36. L. Sutto, S. Marsili, A. Valencia, F. L. Gervasio, From residue coevolution to protein conformational ensembles and functional dynamics. 10.1073/pnas.1508584112 (2015).

37. S. R. Eddy, Accelerated Profile HMM Searches. PLoS Computational Biology 7, e1002195 (2011).

38. M. Ekeberg, C. Lövkvist, Y. Lan, M. Weigt, E. Aurell, Improved contact prediction in proteins: Using pseudolikelihoods to infer Potts models. Physcial Review E 87, 012707 (2013).

39. A. Haldane, R. M. Levy, Influence of multiple-sequence-alignment depth on Potts statistical models of protein covariation. Physcial Review 99, 032405(032415) (2019).

40. Z. G. Gao et al., Identification of essential residues involved in the allosteric modulation of the human A(3) adenosine receptor. Mol Pharmacol 63, 1021–1031 (2003).

41. H. Gutierrez-de-Teran et al., The Role of a Sodium Ion Binding Site in the Allosteric Modulation of the A Adenosine G Protein-Coupled Receptor. Structure 21, 2175–2185 (2013).

42. Y. Miao, A. D. Caliman, J. A. McCammon, Allosteric Effects of Sodium Ion Binding on Activation of the M3 Muscarinic G-Protein Coupled Receptor. Biophysical Journal 108, 1796–1806 (2015).

43. Y. Shang et al., Mechanistic Insights into the Allosteric Modulation of Opioid Receptors by Sodium Ions. Biochemistry 53, 5140–5149 (2014).

44. K. L. White et al., Structural Connection between Activation Microswitch and Allosteric Sodium Site in GPCR Signaling. Structure 26, 259–269 e255 (2018).

45. M. C. Peeters et al., Getting from A to B-exploring the activation motifs of the class B adhesion G protein-coupled receptor subfamily G member 4/GPR112. FASEB J 30, 1836–1848 (2016).

46. O. Fleetwood, P. Matricon, J. Carlsson, L. Delemotte, Energy Landscapes Reveal Agonist Control of G Protein-Coupled Receptor Activation via Microswitches. Biochemistry 59, 880–891 (2020).

47. R. Nygaard, T. M. Frimurer, B. Holst, M. M. Rosenkilde, T. W. Schwartz, Ligand binding and microswitches in 7TM receptor structures. Trends in Pharmacological Sciences 30, 249–259 (2009).

48. S. G. Rasmussen et al., Crystal structure of the β2 adrenergic receptor-Gs protein complex. Nature 477, 549–555 (2011).

49. G. Lebon et al., Agonist-bound adenosine A(2A) receptor structures reveal common features of GPCR activation. Nature 474, 521–U154 (2011).

50. W. Huang et al., Structural insights into micro-opioid receptor activation. Nature 10.1038/nature14886 (2015).

51. A. J. Venkatakrishnan et al., Diverse activation pathways in class A GPCRs converge near the G-protein-coupling region. Nature 536, 484–487 (2016).

52. A. Manglik, A. C. Kruse, Structural Basis for G Protein-Coupled Receptor Activation. Biochemistry 56, 5628–5634 (2017).

53. G. Mattedi, S. Acosta-Gutiérrez, T. Clark, F. L. Grevasio, A combined activation mechanism for the glucagon receptor. Proc Natl Acad Sci USA 117, 15414–15422 (2020).

54. K. J. Gregory, P. M. Sexton, A. Christopoulos, Allosteric modulation of muscarinic acetylcholine receptors. Current neuropharmacology 5, 157–167 (2007).

55. Y. Q. Ping et al., Structures of the glucocorticoid-bound adhesion receptor GPR97-Go complex. Nature 10.1038/s41586-020-03083-w (2021).

56. Z. Rankovic, T. F. Brust, L. M. Bohn, Biased agonism: An emerging paradigm in GPCR drug discovery. Bioorg Med Chem Lett 26, 241–250 (2016).

57. Y. Miao, J. A. McCammon, Graded activation and free energy landscapes of a muscarinic G-protein–coupled receptor. Proc Natl Acad Sci 113, 12162–12167 (2016).

58. L. Seidal, B. Zarzycka, S. A. Zaidi, V. Katritch, I. Coin, Structural insight into the activation of a class B G-protein-coupled receptor by peptide hormones in live human cells. Elife 6 (2017).

59. K. J. Kohlhoff et al., Cloud-based simulations on Google Exacycle reveal ligand modulation of GPCR activation pathways. Nat Chem 6, 15–21 (2014).

60. A. Coucke et al., Direct coevolutionary couplings reflect biophysical residue interactions in proteins. J Chem Phys 145, 174102 (2016).

61. A. Haldane, Levy, R.M., Mi3-GPU: MCMC-based Inversing Ising Inference on GPUs for protein covariation analysis. Computer Physics Communications 107312 (2020).

