## Supplementary material for "Unique Features of Different Classes of G-Protein-Coupled Receptors Revealed from Sequence Coevolutionary and Structural Analysis": SI

### Methods

#### *Sequence datasets*

We built multiple sequence alignments (MSAs) of class A and B GPCRs from the corresponding Pfam IDs of PF00001 and PF00002, respectively. The MSAs included 84,481 sequences for class A and 17,804 sequences for class B of GPCRs (**Table S1**). Any sequences and columns in the MSAs with more than 10% gaps were removed, leaving 37,471 sequences of length 235 residues for class A and 12,645 sequences of length 226 residues for class B of GPCRs. The residue sequences of class A and B GPCRs were also aligned using the *hmmalign* function of HMMER (1) (**Table S2**). A number of these sequences, however, were phylogenetically related and sampled with experimental biases (2-4). One common correction was to downweight similar sequences by assigning a weight $w = {}^{1}/_{n}$ to each sequence, with n being the number of sequences that are more than 40% similar to the target sequence (2-4). The correction resulted in a number of effective sequences $N_{eff} = \sum w$ of 5126 sequences for class A and 902 sequences for class B of GPCRs (**Table S1**).

#### *Sequence coevolutionary Potts model inference*

Potts Hamiltonian models of residue covariation was built based on the assumption that two interacting residues in a protein have correlated mutations during evolution (2, 3). The Potts model seeks to construct a least biased sequence probability distribution $P(S) \propto e^{-H(S)}$ that reproduces one-site $<s_i>$ and two-site $<s_i s_j>$ mutational probabilities of a protein MSA (2, 3, 5). In this equation, *H(S)* is the Hamiltonian and takes the form of $H(S) = \sum_{i=1}^{L} h_i(s_i) + \sum_{i=1}^{L} \sum_{i<j}^{L} J_{ij}(s_i, s_j)$, in which *S* is a sequence of amino acid types (s) at each of L positions, $h_i(s_i)$, or "fields", refers to the single point contribution to the statistical energy of residue $s_i$ at position

i, and $J_{ij}(s_i,s_j)$, or “couplings”, stands for the energy contribution of a position-pair i and j (2, 3, 5). Strong coupling parameters $J_{ij}(s_i,s_j)$ often correspond to direct physical interactions in protein 3D structures (2, 4, 6); therefore, the model is of great interest in the field of structure determination. Mi3-GPU is a newly developed software that solves for real-valued coupling parameters $J_{ij}(s_i,s_j)$ in the Hamiltonian H(S) with few approximations using Markov-Chain Monte-Carlo (MCMC) methods with quasi-Newton optimization (7). Mi3-GPU can be used to construct Potts models with high statistical precision. A full description of the program can be found here (7).

We applied Mi3-GPU to infer the Potts models of class A and B GPCRs. A 21-letter alphabet was used for Potts covariation analysis, including the 20 amino acids plus gap. The damping parameter was set to 0.01 and an l1 regularization of 0.002 throughout the inference. Three rounds of inference were performed for each Potts model. In the first round, $2^{18}$ walkers with 64 MCMC steps were used to reduce the sum-of-square-residuals (SSR), average bivariate marginal relative error (Ferr) and the covariance energy (X). Since our datasets were from Pfam, the X values were not yet fully leveled off after the first round, a second round of inference was thus performed using an increased number of $2^{20}$ walkers with 32-64 MCMC steps to fully level off X values. One final round with $2^{22}$ walkers for four MCMC steps was performed to minimize the finite-sampling error and obtain a model with statistically accurate marginals and negligible residuals (7).

#### ***Residue interaction score using weighted Frobenius norm***

A weighted Frobenius norm (8) was used to obtain a residue pair interaction score from resulting Potts model parameters to control and reduce the contribution of marginals with a large sampling error (3). The interaction score can be calculated from the coupling parameters $J_{\alpha\beta}^{ij}$ using the following formula $I^{ij} = \sqrt{\sum_{\alpha\beta}(w_{\alpha\beta}^{ij} J_{\alpha\beta}^{ij})^2}$ , where $w_{\alpha\beta}^{ij}$ is positive and tunable (3, 7).

***Construction of residue contact frequency maps***

From the Protein Data Bank (PDB) (9, 10) and GPCRdb (11-13), we collected 283 structures of class A GPCRs (94 active, 185 inactive, and four intermediate) and 31 structures of class B GPCRs (16 active, 14 inactive, and one intermediate) (**Table S3**). Among the active GPCR structures, three class A GPCRs were bound by both the agonist and positive allosteric modulator (PAM) (PDBs: 4MQT, 6OIK and 6N48), similarly for one class B GPCR (PDB: 6VCB). Among the inactive GPCR structures, three class A GPCRs were bound by both the antagonist and negative allosteric modulator (NAM) (PDBs: 5X7D, 6OBA and 5T1A), similarly for one class B GPCR (PDB: 5EE7). A summary of the available GPCR structural data is included in **Table S1**.

The contact frequencies between conserved residues of class A and B GPCRs were determined. Two residues were considered in contact if their Cβ-Cβ atom distance was ≤8 Å, unless stated otherwise. For glycine, the Cα atom was used. The residue contact frequency maps were built for classes A and B of GPCRs using all the available structures for comparison with the sequence coevolutionary Potts models. Exceptions of residue contacts that showed significantly high Frobenius norms but low structural contact frequencies were identified in the Potts models of classes A and B GPCRs. Note that a number of these exception residue contacts could result from the arbitrary contact definition since they showed average Cβ-Cβ distances close to the 8Å cutoff (**Table S8**).

Residue contact frequency difference maps were built for active versus inactive GPCR structures to identify residue contacts involved in activation of the two classes of GPCRs. The concepts of switching and repacking contacts for GPCR activation were adopted from Zhou et al. (14). Switching contacts were defined as residue contacts that were present in only one GPCR functional state (active or inactive). Repacking contacts were residue contacts that were present in both GPCR functional states but showed notable changes in contact frequencies. Furthermore, residue contact frequency difference maps were built for active agonist-bound GPCRs in the presence versus absence of PAMs, as well as inactive antagonist-bound GPCRs in the presence versus absence of NAMs, to identify residue contacts tuned by allosteric modulation in the two classes of GPCRs.

**Table S1. Summary of the sequences and experimental PDB structures used for analysis of the class A and B GPCRs**. $N_{init}$ is the initial number of sequences in the multiple sequence alignments (MSAs) obtained from Pfam/Xfam. $N$ and $L$ are the number and length of residue sequences in the MSAs after removal of sequences and columns with more than 10% gaps. $N_{eff}$ is the final number of effective sequences after phylogenetic filtering at a 40% identity threshold. $N_{PDBs}$, $N_{active}$, $N_{inactive}$, $N_{interm}$, $N_{+PAM}$ and $N_{+NAM}$ are the numbers of all PDB structures, active structures, inactive structures, intermediate structures, active with PAM structures and inactive with NAM structures of each GPCR class used to build the residue contact frequency maps.

| Potts Model | Pfam/Xfam | $N_{init}$ | $N$ | $L$ | $N_{eff}$ | $N_{PDBs}$ | $N_{active}$ | $N_{inactive}$ | $N_{interm}$ | $N_{+PAM}$ | $N_{+NAM}$ |
|---|---|---|---|---|---|---|---|---|---|---|---|
| **Class A** | PF00001 | 84481 | 37471 | 235 | 5126 | 283 | 94 | 185 | 4 | 3 | 3 |
| **Class B** | PF00002 | 17804 | 12645 | 226 | 902 | 31 | 16 | 14 | 1 | 1 | 1 |

**Table S2. Sequence alignments of GPCRs: (A) class A (using residue index in β1-adrenoreceptor) and (B) class B (using residue index in glucagon receptor)**. The secondary structures are also shown, with the Ballesteros-Weinstein indexes for helical residues included.

| | | |
|---|---|---|
| **TM1** | | |
| 1 | 1.50 | N59 |
| 2 | 1.51 | V60 |
| 3 | 1.52 | L61 |
| 4 | 1.53 | V62 |
| 5 | 1.54 | I63 |
| 6 | 1.55 | A64 |
| 7 | 1.56 | A65 |
| 8 | 1.57 | I66 |
| 9 | 1.58 | G67 |
| 10 | 1.59 | R68 |
| 11 | 1.60 | T69 |
| **ICL1** | | |
| 12 | | Q70 |
| 13 | | R71 |
| 14 | | L72 |
| 15 | | Q73 |
| **TM2** | | |
| 16 | 2.37 | T74 |
| 17 | 2.38 | L75 |
| 18 | 2.39 | T76 |
| 19 | 2.40 | N77 |
| 20 | 2.41 | L78 |
| 21 | 2.42 | F79 |
| 22 | 2.43 | I80 |
| 23 | 2.44 | T81 |
| 24 | 2.45 | S82 |
| 25 | 2.46 | L83 |
| 26 | 2.47 | A84 |
| 27 | 2.48 | C85 |
| 28 | 2.49 | A86 |
| 29 | 2.50 | D87 |
| 30 | 2.51 | L88 |
| 31 | 2.52 | V89 |
| 32 | 2.53 | M90 |
| 33 | 2.55 | L92 |
| 34 | 2.56 | L93 |
| 35 | 2.57 | V94 |
| 36 | 2.58 | V95 |
| 37 | 2.59 | P96 |
| 38 | 2.60 | F97 |
| 39 | 2.61 | G98 |
| 40 | 2.62 | A99 |
| 41 | 2.63 | T100 |
| 42 | 2.64 | L101 |
| 43 | 2.65 | V102 |
| 44 | 2.67 | R104 |
| **ECL1** | | |
| 45 | | G105 |
| 46 | | T106 |
| 47 | | W107 |
| 48 | | L108 |
| 49 | | W109 |
| **TM3** | | |
| 50 | 3.21 | G110 |
| 51 | 3.22 | S111 |
| 52 | 3.23 | F112 |
| 53 | 3.24 | L113 |
| 54 | 3.25 | C114 |
| 55 | 3.26 | E115 |
| 56 | 3.27 | C116 |
| 57 | 3.28 | W117 |
| 58 | 3.29 | T118 |
| 59 | 3.30 | S119 |
| 60 | 3.31 | L120 |
| 61 | 3.32 | D121 |
| 62 | 3.33 | V122 |
| 63 | 3.34 | L123 |
| 64 | 3.35 | C124 |
| 65 | 3.36 | V125 |
| 66 | 3.37 | T126 |
| 67 | 3.38 | A127 |
| 68 | 3.39 | S128 |
| 69 | 3.40 | I129 |
| 70 | 3.41 | E130 |
| 71 | 3.42 | T131 |
| 72 | 3.43 | L132 |
| 73 | 3.44 | C133 |
| 74 | 3.45 | V134 |
| 75 | 3.46 | I135 |
| 76 | 3.47 | A136 |
| 77 | 3.48 | I137 |
| 78 | 3.49 | D138 |
| 79 | 3.50 | R139 |
| 80 | 3.51 | Y140 |
| 81 | 3.52 | L141 |
| 82 | 3.53 | A142 |
| 83 | 3.54 | I143 |
| 84 | 3.55 | T144 |
| 85 | 3.56 | S145 |
| **ICL2** | | |
| 86 | | P146 |
| 87 | | F147 |
| 88 | | R148 |
| 89 | | Y149 |
| 90 | | Q150 |
| 91 | | S151 |
| 92 | | L152 |
| 93 | | M153 |
| **TM4** | | |
| 94 | 4.38 | T154 |
| 95 | 4.39 | R155 |
| 96 | 4.40 | A156 |
| 97 | 4.41 | R157 |
| 98 | 4.42 | A158 |
| 99 | 4.43 | K159 |
| 100 | 4.44 | V160 |
| 101 | 4.45 | I161 |
| 102 | 4.46 | I162 |
| 103 | 4.47 | C163 |
| 104 | 4.48 | T164 |
| 105 | 4.49 | V165 |
| 106 | 4.50 | W166 |
| 107 | 4.51 | A167 |
| 108 | 4.52 | I168 |
| 109 | 4.53 | S169 |
| 110 | 4.54 | A170 |
| 111 | 4.55 | L171 |
| 112 | 4.56 | V172 |
| 113 | 4.57 | S173 |
| 114 | 4.58 | F174 |
| 115 | 4.59 | L175 |
| 116 | 4.60 | P176 |
| 117 | 4.61 | I177 |
| 118 | 4.62 | M178 |
| 119 | 4.63 | M179 |
| 120 | 4.64 | H180 |
| **ECL2** | | |
| 121 | | W181 |
| 122 | | W182 |
| 123 | | R183 |
| 124 | | D184 |
| 125 | | E185 |
| 126 | | D186 |
| 127 | | P187 |
| 128 | | Q188 |
| 129 | | G197 |
| 130 | | C198 |
| 131 | | C199 |
| 132 | | D200 |
| 133 | | F201 |
| 134 | | V202 |
| 135 | | T203 |
| **TM5** | | |
| 136 | 5.35 | N204 |
| 137 | 5.36 | R205 |
| 138 | 5.37 | A206 |
| 139 | 5.38 | Y207 |
| 140 | 5.39 | A208 |
| 141 | 5.40 | I209 |
| 142 | 5.41 | A210 |
| 143 | 5.42 | S211 |
| 144 | 5.43 | S212 |
| 145 | 5.44 | I213 |
| 146 | 5.45 | I214 |
| 147 | 5.46 | S215 |
| 148 | 5.47 | F216 |
| 149 | 5.48 | Y217 |
| 150 | 5.49 | I218 |
| 151 | 5.50 | P219 |
| 152 | 5.51 | L220 |
| 153 | 5.52 | L221 |
| 154 | 5.53 | I222 |
| 155 | 5.54 | M223 |
| 156 | 5.55 | I224 |
| 157 | 5.56 | F225 |
| 158 | 5.57 | V226 |
| 159 | 5.58 | Y227 |
| 160 | 5.59 | L228 |
| 161 | 5.60 | R229 |
| 162 | 5.61 | V230 |
| 163 | 5.62 | Y231 |
| 164 | 5.63 | R232 |
| 165 | 5.64 | E233 |
| 166 | 5.65 | A234 |
| 167 | 5.66 | K235 |
| 168 | 5.67 | E236 |
| 169 | 5.68 | Q237 |
| 170 | 5.69 | I238 |
| 171 | 5.70 | R239 |
| 172 | 5.71 | K240 |
| 173 | 5.72 | I241 |
| 174 | 5.73 | D242 |
| 175 | 5.74 | R243 |
| **TM6** | | |
| 176 | 6.29 | R284 |
| 177 | 6.30 | E285 |
| 178 | 6.31 | H286 |
| 179 | 6.32 | K287 |
| 180 | 6.33 | A288 |
| 181 | 6.34 | L289 |
| 182 | 6.35 | K290 |
| 183 | 6.36 | T291 |
| 184 | 6.37 | L292 |
| 185 | 6.38 | G293 |
| 186 | 6.39 | I294 |
| 187 | 6.40 | I295 |
| 188 | 6.41 | M296 |
| 189 | 6.42 | G297 |
| 190 | 6.43 | V298 |
| 191 | 6.44 | F299 |
| 192 | 6.45 | T300 |
| 193 | 6.46 | L301 |
| 194 | 6.47 | C302 |
| 195 | 6.48 | W303 |
| 196 | 6.49 | L304 |
| 197 | 6.50 | P305 |
| 198 | 6.51 | F306 |
| 199 | 6.52 | F307 |
| 200 | 6.53 | L308 |
| 201 | 6.54 | V309 |
| 202 | 6.55 | N310 |
| 203 | 6.56 | I311 |
| 204 | 6.57 | V312 |
| 205 | 6.58 | N313 |
| 206 | 6.59 | V314 |
| 207 | 6.60 | F315 |
| 208 | 6.61 | N316 |
| **ECL3** | | |
| 209 | | R317 |
| 210 | | D318 |
| 211 | | L319 |
| 212 | | V320 |
| **TM7** | | |
| 213 | 7.31 | P321 |
| 214 | 7.32 | D322 |
| 215 | 7.33 | W323 |
| 216 | 7.34 | L324 |
| 217 | 7.35 | F325 |
| 218 | 7.36 | V326 |
| 219 | 7.37 | F327 |
| 220 | 7.38 | F328 |
| 221 | 7.39 | N329 |
| 222 | 7.40 | W330 |
| 223 | 7.41 | L331 |
| 224 | 7.42 | G332 |
| 225 | 7.43 | Y333 |
| 226 | 7.44 | A334 |
| 227 | 7.45 | N335 |
| 228 | 7.46 | S336 |
| 229 | 7.47 | A337 |
| 230 | 7.48 | F338 |
| 231 | 7.49 | N339 |
| 232 | 7.50 | P340 |
| 233 | 7.51 | I341 |
| 234 | 7.52 | I342 |
| 235 | 7.53 | Y343 |

**(A) Class A**

| | | |
|---|---|---|
| **TM1** | | |
| 1 | 1.36 | Q142 |
| 2 | 1.37 | V143 |
| 3 | 1.38 | M144 |
| 4 | 1.39 | Y145 |
| 5 | 1.40 | T146 |
| 6 | 1.41 | V147 |
| 7 | 1.42 | G148 |
| 8 | 1.43 | Y149 |
| 9 | 1.44 | S150 |
| 10 | 1.45 | L151 |
| 11 | 1.46 | S152 |
| 12 | 1.47 | L153 |
| 13 | 1.48 | G154 |
| 14 | 1.49 | A155 |
| 15 | 1.50 | L156 |
| 16 | 1.51 | L157 |
| 17 | 1.52 | L158 |
| 18 | 1.53 | A159 |
| 19 | 1.54 | L160 |
| 20 | 1.55 | A161 |
| 21 | 1.56 | I162 |
| 22 | 1.57 | L163 |
| 23 | 1.58 | G164 |
| 24 | 1.59 | G165 |
| 25 | 1.60 | L166 |
| **ICL1** | | |
| 26 | | S167 |
| 27 | | K168 |
| 28 | | L169 |
| 29 | | H170 |
| **TM2** | | |
| 30 | 2.37 | C171 |
| 31 | 2.38 | T172 |
| 32 | 2.39 | R173 |
| 33 | 2.40 | N174 |
| 34 | 2.41 | A175 |
| 35 | 2.42 | I176 |
| 36 | 2.43 | H177 |
| 37 | 2.44 | A178 |
| 38 | 2.45 | N179 |
| 39 | 2.46 | L180 |
| 40 | 2.47 | F181 |
| 41 | 2.48 | A182 |
| 42 | 2.49 | S183 |
| 43 | 2.50 | F184 |
| 44 | 2.51 | V185 |
| 45 | 2.52 | L186 |
| 46 | 2.53 | K187 |
| 47 | 2.54 | A188 |
| 48 | 2.55 | S189 |
| 49 | 2.56 | S190 |
| 50 | 2.57 | V191 |
| 51 | 2.58 | L192 |
| 52 | 2.59 | V193 |
| 53 | 2.60 | I194 |
| 54 | 2.61 | D195 |
| 55 | 2.62 | G196 |
| 56 | 2.63 | L197 |
| 57 | 2.64 | L198 |
| 58 | 2.65 | R199 |
| 59 | 2.66 | T200 |
| 60 | 2.67 | R201 |
| **TM3** | | |
| 61 | 3.23 | A222 |
| 62 | 3.24 | G223 |
| 63 | 3.25 | C224 |
| 64 | 3.26 | R225 |
| 65 | 3.27 | V226 |
| 66 | 3.28 | A227 |
| 67 | 3.29 | A228 |
| 68 | 3.30 | V229 |
| 69 | 3.31 | F230 |
| 70 | 3.32 | M231 |
| 71 | 3.33 | Q232 |
| 72 | 3.34 | Y233 |
| 73 | 3.35 | G234 |
| 74 | 3.36 | I235 |
| 75 | 3.37 | V236 |
| 76 | 3.38 | A237 |
| 77 | 3.39 | N238 |
| 78 | 3.40 | Y239 |
| 79 | 3.41 | C240 |
| 80 | 3.42 | W241 |
| 81 | 3.43 | L242 |
| 82 | 3.44 | L243 |
| 83 | 3.45 | V244 |
| 84 | 3.46 | E245 |
| 85 | 3.47 | G246 |
| 86 | 3.48 | L247 |
| 87 | 3.49 | Y248 |
| 88 | 3.50 | L249 |
| 89 | 3.51 | H250 |
| 90 | 3.52 | N251 |
| 91 | 3.53 | L252 |
| 92 | 3.54 | L253 |
| 93 | 3.55 | G254 |
| 94 | 3.56 | L255 |
| **ICL2** | | |
| 95 | | A256 |
| 96 | | T257 |
| 97 | | L258 |
| 98 | | P259 |
| 99 | | E260 |
| 100 | | R261 |
| **TM4** | | |
| 101 | 4.40 | S262 |
| 102 | 4.41 | F263 |
| 103 | 4.42 | F264 |
| 104 | 4.43 | S265 |
| 105 | 4.44 | L266 |
| 106 | 4.45 | Y267 |
| 107 | 4.46 | L268 |
| 108 | 4.47 | G269 |
| 109 | 4.48 | I270 |
| 110 | 4.49 | G271 |
| 111 | 4.50 | W272 |
| 112 | 4.51 | G273 |
| 113 | 4.52 | A274 |
| 114 | 4.53 | P275 |
| 115 | 4.54 | M276 |
| 116 | 4.55 | L277 |
| 117 | 4.56 | F278 |
| 118 | 4.57 | V279 |
| 119 | 4.58 | V280 |
| 120 | 4.59 | P281 |
| 121 | 4.60 | W282 |
| 122 | 4.61 | A283 |
| 123 | 4.62 | V284 |
| 124 | 4.63 | V285 |
| 125 | 4.64 | K286 |
| 126 | 4.65 | C287 |
| 127 | 4.66 | L288 |
| 128 | 4.67 | F289 |
| 129 | 4.68 | E290 |
| **ECL2** | | |
| 130 | | N291 |
| 131 | | V292 |
| 132 | | Q293 |
| 133 | | C294 |
| 134 | | W295 |
| 135 | | T296 |
| 136 | | S297 |
| 137 | | N298 |
| **TM5** | | |
| 138 | 5.36 | D299 |
| 139 | 5.37 | N300 |
| 140 | 5.38 | G302 |
| 141 | 5.39 | F303 |
| 142 | 5.40 | W304 |
| 143 | 5.41 | W305 |
| 144 | 5.42 | I306 |
| 145 | 5.43 | L307 |
| 146 | 5.44 | R308 |
| 147 | 5.45 | F309 |
| 148 | 5.46 | P310 |
| 149 | 5.47 | V311 |
| 150 | 5.48 | F312 |
| 151 | 5.49 | L313 |
| 152 | 5.50 | A314 |
| 153 | 5.51 | I315 |
| 154 | 5.52 | L316 |
| 155 | 5.53 | I317 |
| 156 | 5.54 | N318 |
| 157 | 5.55 | F319 |
| 158 | 5.56 | F320 |
| 159 | 5.57 | I321 |
| 160 | 5.58 | F322 |
| 161 | 5.59 | V323 |
| 162 | 5.60 | R324 |
| 163 | 5.61 | I325 |
| 164 | 5.62 | V326 |
| 165 | 5.63 | Q327 |
| 166 | 5.64 | L328 |
| 167 | 5.65 | L329 |
| 168 | 5.66 | V330 |
| 169 | 5.67 | A331 |
| 170 | 5.68 | K332 |
| 171 | 5.69 | L333 |
| 172 | 5.70 | R334 |
| 173 | 5.71 | A335 |
| **ICL3** | | |
| 174 | | R336 |
| 175 | | Q337 |
| 176 | | M338 |
| 177 | | H339 |
| 178 | | H340 |
| 179 | | T341 |
| **TM6** | | |
| 180 | 6.28 | D342 |
| 181 | 6.29 | Y343 |
| 182 | 6.30 | K344 |
| 183 | 6.31 | F345 |
| 184 | 6.32 | R346 |
| 185 | 6.33 | L347 |
| 186 | 6.34 | A348 |
| 187 | 6.35 | K349 |
| 188 | 6.36 | S350 |
| 189 | 6.37 | T351 |
| 190 | 6.38 | L352 |
| 191 | 6.39 | T353 |
| 192 | 6.40 | L354 |
| 193 | 6.41 | I355 |
| 194 | 6.42 | P356 |
| 195 | 6.43 | L357 |
| 196 | 6.44 | L358 |
| 197 | 6.45 | G359 |
| 198 | 6.46 | V360 |
| 199 | 6.47 | H361 |
| 200 | 6.48 | E362 |
| 201 | 6.49 | V363 |
| 202 | 6.50 | V364 |
| 203 | 6.51 | F365 |
| 204 | 6.52 | A366 |
| 205 | 6.53 | F367 |
| **ECL3** | | |
| 206 | | V368 |
| 207 | | T369 |
| 208 | | D370 |
| 209 | | E371 |
| 210 | | H372 |
| 211 | | A373 |
| **TM7** | | |
| 212 | 7.34 | K381 |
| 213 | 7.35 | L382 |
| 214 | 7.36 | F383 |
| 215 | 7.37 | F384 |
| 216 | 7.38 | D385 |
| 217 | 7.39 | L386 |
| 218 | 7.40 | F387 |
| 219 | 7.41 | L388 |
| 220 | 7.42 | S389 |
| 221 | 7.43 | S390 |
| 222 | 7.44 | F391 |
| 223 | 7.45 | Q392 |
| 224 | 7.46 | G393 |
| 225 | 7.47 | L394 |
| 226 | 7.48 | L395 |

### (B) Class B

**Table S3. GPCR structures used for construction of residue contact frequency maps: (A) class A and (B) class B.** For class A GPCRs, there are 94 active, 185 inactive and 4 intermediate PDB structures. For class B GPCRs, there are 16 active, 14 inactive and 1 intermediate PDB structures. The active+PAM vs -PAM are active agonist-bound structures with and without PAMs used to calculate the differences in residue contact frequencies upon PAM binding. The inactive+NAM vs -NAM are inactive antagonist-bound structures with and without NAMs used to calculate the differences in residue contact frequencies upon NAM binding.

| Active (94) | | Inactive (185) | | | Intermediate (4) | Active+PAM (3) | Inactive+NAM (3) |
|---|---|---|---|---|---|---|---|
| 2X72 | 6AK3 | 1GZM | 5A8E | 5ZTY | 6DS0 | 4MQT | 5X7D |
| 2YDO | 6B73 | 1U19 | 5D6L | 6A93 | 6LI0 | 6OIK | 6OBA |
| 2YDV | 6CMO | 2G87 | 5DHG | 6A94 | 6LI1 | 6N48 | 5T1A |
| 3CAP | 6D9H | 2HPY | 5DHH | 6AKY | 6LI2 | | |
| 3DQB | 6DDE | 2I35 | 5F8U | 6AQF | | **-PAM (3)** | **-NAM (3)** |
| 3P0G | 6DDF | 2I36 | 5GLH | 6CM4 | | 4MQS | 6PRZ |
| 3PQR | 6DO1 | 2J4Y | 5GLI | 6D27 | | 6U1N | 6PS0 |
| 3PXO | 6E67 | 2PED | 5IU4 | 6GPS | | 6NI3 | 6GPS |
| 3QAK | 6FK6 | 2RH1 | 5IU7 | 6GPX | | | |
| 3SN6 | 6FK7 | 2Y00 | 5IU8 | 6GT3 | | | |
| 4A4M | 6FK8 | 2Y01 | 5IUA | 6HLL | | | |
| 4BEY | 6FK9 | 2Y02 | 5IUB | 6HLO | | | |
| 4BEZ | 6FKA | 2Y03 | 5JQH | 6HLP | | | |
| 4GRV | 6FKB | 2Y04 | 5JTB | 6IGK | | | |
| 4J4Q | 6FKC | 2YCW | 5K2A | 6IGL | | | |
| 4LDE | 6FKD | 2Z73 | 5K2B | 6J20 | | | |
| 4LDL | 6FUF | 2ZIY | 5K2C | 6J21 | | | |
| 4LDO | 6GDG | 3C9L | 5K2D | 6JZH | | | |
| 4MQS | 6H7J | 3C9M | 5MZJ | 6K1Q | | | |
| 4MQT | 6H7L | 3D4S | 5MZP | 6KNM | | | |
| 4PXF | 6H7M | 3EML | 5N2R | 6KPC | | | |
| 4QKX | 6H7N | 3NY8 | 5N2S | 6KQI | | | |
| 4UHR | 6H7O | 3OAX | 5NLX | 6LRY | | | |
| 4X1H | 6IBL | 3ODU | 5NM2 | 6LUQ | | | |
| 4XEE | 6JOD | 3PBL | 5NM4 | 6ME2 | | | |
| 4XES | 6KPF | 3PDS | 5O9H | 6ME3 | | | |
| 4XT1 | 6LI3 | 3PWH | 5OLG | 6ME5 | | | |
| 4XT3 | 6LW5 | 3REY | 5OLH | 6ME6 | | | |
| 4ZWJ | 6M9T | 3RFM | 5OLO | 6ME7 | | | |
| 5C1M | 6MXT | 3RZE | 5OLV | 6ME8 | | | |
| 5DGY | 6N48 | 3UON | 5OLZ | 6ME9 | | | |
| 5DYS | 6N4B | 3UZA | 5OM1 | 6MEO | | | |
| 5EN0 | 6NI3 | 3UZC | 5OM4 | 6MET | | | |
| 5G53 | 6OIJ | 3VG9 | 5T1A | 6MH8 | | | |
| 5T04 | 6OIK | 3VGA | 5TE5 | 6OBA | | | |
| 5TE3 | 6OMM | 3ZPQ | 5TGZ | 6OL9 | | | |
| 5TUD | 6OS0 | 3ZPR | 5U09 | 6PRZ | | | |
| 5UNF | 6OS1 | 4AMI | 5UEN | 6PS0 | | | |
| 5UNG | 6OS2 | 4AMJ | 5UIG | 6PS1 | | | |
| 5UNH | 6OS9 | 4BVN | 5UIW | 6PS3 | | | |
| 5W0P | 6OSA | 4DJH | 5UVI | 6PS4 | | | |
| 5WB1 | 6OY9 | 4DKL | 5V54 | 6PS5 | | | |
| 5WB2 | 6OYA | 4EA3 | 5VBL | 6PS6 | | | |
| 5WF5 | 6PWC | 4EIY | 5VRA | 6PS7 | | | |
| 5WKT | 6QNO | 4EJ4 | 5WIU | 6PS8 | | | |
| 5XJM | 6U1N | 4GBR | 5WIV | 6QZH | | | |
| 5XRA | 6UP7 | 4GPO | 5WQC | 6TO7 | | | |
| | | 4IAQ | 5WS3 | 6TOD | | | |
| | | 4IAR | 5X7D | 6TOS | | | |
| | | 4MBS | 5X93 | 6TOT | | | |
| | | 4N6H | 5XCV | 6TP3 | | | |
| | | 4RWD | 5XPR | 6TP4 | | | |
| | | 4S0V | 5YC8 | 6TP6 | | | |
| | | 4U15 | 5YHL | 6TPG | | | |
| | | 4U16 | 5YWY | 6TPJ | | | |
| | | 4WW3 | 5ZBH | 6TPN | | | |
| | | 4YAY | 5ZBQ | 6TQ4 | | | |
| | | 4Z34 | 5ZHP | 6TQ6 | | | |
| | | 4Z35 | 5ZK3 | 6TQ7 | | | |
| | | 4Z36 | 5ZK8 | 6TQ9 | | | |
| | | 4ZJC | 5ZKB | 6VI4 | | | |
| | | 4ZUD | 5ZKC | | | | |

**(A) Class A**

| **Active (16)** | **Inactive (14)** | **Intermediate (1)** | **Active+PAM (1)** | **Inactive+NAM (1)** |
|---|---|---|---|---|
| 5UZ7 | 4K5Y | 5NX2 | 6VCB | 5EE7 |
| 5VAI | 4L6R | | | |
| 6B3J | 4Z9G | | | |
| 6E3Y | 5EE7 | | | |
| 6LMK | 5VEW | | **-PAM (1)** | **-NAM (1)** |
| 6LML | 5VEX | | 6B3J | 5XEZ |
| 6LPB | 5XEZ | | | |
| 6NBF | 5XF1 | | | |
| 6NBH | 5YQZ | | | |
| 6NBI | 6FJ3 | | | |
| 6NIY | 6KJV | | | |
| 6P9Y | 6KK1 | | | |
| 6PB1 | 6KK7 | | | |
| 6UUN | 6LN2 | | | |
| 6UUS | | | | |
| 6UVA | | | | |

**(B) Class B**

**Table S4. Top ranked exception residue contacts predicted in the Potts model that are important for activation and allosteric modulation of class A GPCRs**. Only residue pairs with at least five residues apart in the numbering scheme were considered. The amino acid identities are listed using the $\beta_1$-adrenoreceptor as a model class A GPCR. *FB* is the Frobenius norms calculated from the MSA by Potts model. $CF_{PDBs}$, $CF_{Active}$ and $CF_{Inactive}$ are the contact frequencies in all PDB, active and inactive class A GPCR structures. $\Delta CF_{PAM}$ is the difference in contact frequencies between agonist-PAM and agonist bound GPCR structures and $\Delta CF_{NAM}$ is the difference in contact frequencies between antagonist-NAM and antagonist bound GPCR structures. Exception residue contacts that are explained by GPCR activation and allosteric are marked "Activation", "Inactivation", and "Allostery" in Role. Other unexplained exception residue contacts which have average – standard deviation of distance ≤9.0Å in all PDB structures in **Table S8A** are marked "Contact Definition" in Role.

| Rank | Residues | FB | $CF_{PDBs}$ | $CF_{Active}$ | $CF_{Inactive}$ | $\Delta CF_{PAM}$ | $\Delta CF_{NAM}$ | Role |
|---|---|---|---|---|---|---|---|---|
| 4 | Y3.51-S3.56 | 0.33 | 0.11 | 0.14 | 0.10 | 0 | -0.33 | Allostery |
| 11 | A2.49-W4.50 | 0.23 | 0.03 | 0.06 | 0.01 | -0.67 | 0 | Allostery |
| 14 | C3.44-V5.57 | 0.21 | 0.28 | 0.10 | 0.38 | 0 | 0 | Inactivation |
| 18 | W4.50-P7.50 | 0.20 | 0 | 0 | 0 | 0 | 0 | - |
| 20 | $C198^{ECL2}$-N5.35 | 0.20 | 0.04 | 0.01 | 0.05 | 0 | 0 | - |
| 28 | Y3.51-F5.56 | 0.19 | 0.20 | 0.38 | 0.12 | 0 | 0 | Activation |
| 29 | T3.42-W4.50 | 0.19 | 0.15 | 0.11 | 0.18 | 0 | 0 | Contact Definition |
| 30 | D3.49-$Y149^{ICL2}$ | 0.19 | 0.04 | 0.06 | 0.03 | -0.33 | 0 | Allostery |
| 34 | S3.30-W6.48 | 0.18 | 0 | 0 | 0 | 0 | 0 | - |
| 37 | R3.50-N7.45 | 0.18 | 0 | 0 | 0 | 0 | 0 | - |
| 39 | $C198^{ECL2}$-$T203^{ECL2}$ | 0.17 | 0.03 | 0.01 | 0.04 | 0 | 0 | - |
| 41 | D3.32-Y7.43 | 0.17 | 0.08 | 0.05 | 0.09 | 0 | 0 | Contact Definition |
| 44 | Y5.58-P7.50 | 0.17 | 0 | 0 | 0 | 0 | 0 | - |
| 48 | F2.42-$M153^{ICL2}$ | 0.17 | 0 | 0 | 0.01 | 0 | 0 | - |
| 53 | N6.61-D7.32 | 0.16 | 0.01 | 0 | 0.02 | 0 | 0 | - |
| 56 | L4.59-Y5.38 | 0.16 | 0.07 | 0.07 | 0.06 | 0 | 0 | - |
| 57 | I6.40-P6.50 | 0.16 | 0 | 0 | 0 | 0 | 0 | - |
| 60 | F5.47-L6.49 | 0.16 | 0.26 | 0.49 | 0.15 | 0.33 | 0 | Activation |
| 61 | L2.46-S7.46 | 0.16 | 0 | 0 | 0.01 | 0 | 0 | Contact Definition |
| 62 | S4.53-N7.49 | 0.16 | 0 | 0 | 0 | 0 | 0 | - |

**Table S5. Residue contacts that are important for (A) activation and (B) allosteric modulation of class A GPCRs. (A)** The amino acid identities are listed using the $\beta_1$-adrenoreceptor as a model class A GPCR. $CF_{PDBs}$, $CF_{Active}$ and $CF_{Inactive}$ are the contact frequencies in all PDB, active and inactive structures of class A GPCRs. $\Delta CF_{Activate}$ is the difference in contact frequencies between the active and inactive GPCR structures. **(B)** The amino acid identities are taken from the M2 receptor (PAM) and $\beta_2$-adrenoreceptor (NAM). $CF_{PAM}$, $CF'_{Active}$, $CF_{NAM}$ and $CF'_{Inactive}$ are the contact frequencies in class A GPCR structures bound by the agonist and PAM (PDBs: 4MQT, 6OIK, 6N48), only agonist (PDBs: 4MQS, 6U1N, 6NI3), antagonist and NAM (PDBs: 5X7D, 6OBA, 5T1A), and only antagonist (PDBs: 6PRZ, 6PS0, 6GPS). $\Delta CF_{PAM}$ is the difference in contact frequencies between agonist-PAM and agonist bound GPCR structures, and $\Delta CF_{NAM}$ is the difference in contact frequencies between antagonist-NAM and antagonist bound GPCR structures.

| Rank | Residues | $CF_{PDBs}$ | $CF_{Active}$ | $CF_{Inactive}$ | $\Delta CF_{Activate}$ | Type |
|---|---|---|---|---|---|---|
| 1 | A5.65-A6.33 | 0.28 | 0.84 | 0 | 0.84 | *Switching* |
| 2 | L2.46-Y7.53 | 0.29 | 0.84 | 0.02 | 0.82 | Repacking |
| 3 | L3.43-N7.49 | 0.28 | 0.82 | 0.01 | 0.81 | Repacking |
| 4 | V1.53-Y7.53 | 0.66 | 0.14 | 0.93 | -0.79 | Repacking |
| 5 | S3.39-S7.46 | 0.37 | 0.87 | 0.11 | 0.76 | Repacking |
| 6 | S3.39-N7.49 | 0.35 | 0.85 | 0.09 | 0.76 | Repacking |
| 7 | I3.46-Y7.53 | 0.25 | 0.76 | 0 | 0.76 | *Switching* |
| 8 | V1.53-N7.49 | 0.51 | 0.01 | 0.76 | -0.75 | Repacking |
| 9 | M2.53-S3.39 | 0.35 | 0.83 | 0.1 | 0.73 | Repacking |
| 10 | M5.54-I6.40 | 0.25 | 0.70 | 0.01 | 0.69 | Repacking |
| 11 | I6.40-N7.49 | 0.54 | 0.10 | 0.79 | -0.69 | Repacking |
| 12 | A3.47-L6.37 | 0.51 | 0.09 | 0.74 | -0.65 | Repacking |
| 13 | I3.46-L6.37 | 0.42 | 0 | 0.64 | -0.64 | *Switching* |
| 14 | L3.43-M6.41 | 0.44 | 0.02 | 0.65 | -0.63 | Repacking |
| 15 | R3.50-L6.34 | 0.42 | 0 | 0.63 | -0.63 | *Switching* |
| 16 | T6.36-I7.52 | 0.61 | 0.21 | 0.83 | -0.62 | Repacking |
| 17 | Y5.58-I6.40 | 0.20 | 0.61 | 0 | 0.61 | *Switching* |
| 18 | D2.50-P7.50 | 0.52 | 0.91 | 0.31 | 0.60 | Repacking |
| 19 | M5.54-F6.44 | 0.22 | 0.62 | 0.02 | 0.60 | Repacking |
| 20 | T6.36-Y7.53 | 0.48 | 0.09 | 0.69 | -0.60 | Repacking |
| 21 | V5.61-L6.34 | 0.51 | 0.12 | 0.72 | -0.60 | Repacking |
| 22 | L6.46-L7.41 | 0.37 | 0.77 | 0.18 | 0.59 | Repacking |
| 23 | A3.47-M6.41 | 0.43 | 0.04 | 0.63 | -0.59 | Repacking |
| 24 | S3.39-F6.44 | 0.37 | 0 | 0.56 | -0.56 | *Switching* |
| 25 | Y5.62-L6.37 | 0.18 | 0.54 | 0 | 0.54 | *Switching* |
| 26 | A2.47-N7.49 | 0.75 | 0.40 | 0.93 | -0.53 | Repacking |
| 27 | V5.61-A6.33 | 0.20 | 0.54 | 0.02 | 0.52 | Repacking |
| 28 | V5.61-L6.37 | 0.25 | 0.60 | 0.08 | 0.52 | Repacking |
| 29 | R3.50-L6.37 | 0.44 | 0.11 | 0.63 | -0.52 | Repacking |
| 30 | A5.65-H6.31 | 0.41 | 0.07 | 0.58 | -0.51 | Repacking |

**(A)**

| Rank | Residues | $CF_{PAM}$ | $CF'_{Active}$ | $\Delta CF_{PAM}$ | Modulator |
|---|---|---|---|---|---|
| 1 | L3.27-L4.62 | 1.00 | 0 | 1.00 | PAM |
| 2 | A6.43-N7.49 | 0 | 1.00 | -1.00 | PAM |
| 3 | V3.34-L4.56 | 0.67 | 0 | 0.67 | PAM |
| 4 | V3.40-S4.53 | 0 | 0.67 | -0.67 | PAM |
| 5 | D3.49-A4.42 | 0.67 | 0 | 0.67 | PAM |
| 6 | N2.39-R3.50 | 1.00 | 0.33 | 0.67 | PAM |
| 7 | N2.39-C3.53 | 1.00 | 0.33 | 0.67 | PAM |
| 8 | F3.52-T34.52 | 0 | 0.67 | -0.67 | PAM |
| 9 | A2.49-W4.50 | 0 | 0.67 | -0.67 | PAM |
| 10 | L23.52-C45.50 | 0 | 0.67 | -0.67 | PAM |
| 11 | R3.50-L5.57 | 0 | 0.67 | -0.67 | PAM |
| 12 | S4.53-A5.46 | 0 | 0.67 | -0.67 | PAM |
| 13 | V5.51-W6.48 | 1.00 | 0.33 | 0.67 | PAM |
| **Rank** | **Residues** | $CF_{NAM}$ | $CF'_{Inactive}$ | $\Delta CF_{NAM}$ | **Modulator** |
| 1 | T6.43-N7.49 | 0 | 1 | -1.00 | NAM |
| 2 | N6.55-Y7.35 | 0.33 | 1 | -0.67 | NAM |
| 3 | A2.49-V3.36 | 1.00 | 0.33 | 0.67 | NAM |
| 4 | I3.31-S4.57 | 0.67 | 0 | 0.67 | NAM |
| 5 | T2.37-K4.39 | 0.67 | 0 | 0.67 | NAM |
| 6 | F2.42-I4.45 | 0 | 0.67 | -0.67 | NAM |
| 7 | T2.39-A6.33 | 0.33 | 1 | -0.67 | NAM |
| 8 | I2.43-I6.40 | 0 | 0.67 | -0.67 | NAM |
| 9 | R3.50-V5.57 | 0 | 0.67 | -0.67 | NAM |
| 10 | S4.57-S5.42 | 0 | 0.67 | -0.67 | NAM |
| 11 | A5.39-H6.58 | 0.67 | 0 | 0.67 | NAM |
| 12 | V6.59-D300$^{ECL3}$ | 1.00 | 0.33 | 0.67 | NAM |

**(B)**

**Table S6. Top ranked exception residue contacts predicted in the Potts model that are important for activation of class B GPCRs**. Only residue pairs with at least five residues apart were considered. The amino acid identities are listed using the glucagon receptor as a model class B GPCR. *FB* is the Frobenius norms calculated from the MSA by Potts model. $CF_{PDBs}$, $CF_{Active}$ and $CF_{Inactive}$ are the contact frequencies in all PDB, active and inactive class B GPCR structures. $\Delta CF_{PAM}$ is the difference in contact frequencies between agonist-PAM and agonist bound GPCR structures and $\Delta CF_{NAM}$ is the difference in contact frequencies between antagonist-NAM and antagonist bound GPCR structures. Because the available structures of class B GPCRs all function as secretin receptors, residue contacts are either present in most of the structures or none of the structures. Exception residue contacts that are explained by GPCR activation and allosteric are marked "Activation", "Inactivation", and "Allostery" in Role. Other unexplained exception residue contacts which have average – standard deviation of distance ≤9.0Å in all PDB structures in **Table S8B** are marked "Contact Definition" in Role.

| Rank | Residues | FB | $CF_{PDBs}$ | $CF_{Active}$ | $CF_{Inactive}$ | $\Delta CF_{PAM}$ | $\Delta CF_{NAM}$ | Role |
|---|---|---|---|---|---|---|---|---|
| 2 | S2.49-W4.50 | 0.36 | 0.23 | 0.31 | 0.14 | 0 | 0 | Activation |
| 7 | Q3.33-L4.66 | 0.29 | 0 | 0 | 0 | 0 | 0 | - |
| 19 | W295$^{ECL2}$-L5.43 | 0.25 | 0 | 0 | 0 | 0 | 0 | - |
| 21 | H2.43-L6.40 | 0.25 | 0 | 0 | 0 | 0 | 0 | - |
| 23 | L6.44-S7.42 | 0.24 | 0 | 0 | 0 | 0 | 0 | - |
| 27 | Q3.33-E6.48 | 0.24 | 0 | 0 | 0 | 0 | 0 | - |
| 30 | F5.58-L6.33 | 0.23 | 0 | 0 | 0 | 0 | 0 | - |
| 32 | N5.54-L6.43 | 0.22 | 0 | 0 | 0 | 0 | 0 | - |
| 33 | A2.54-L7.39 | 0.22 | 0 | 0 | 0 | 0 | 0 | - |
| 35 | K2.53-L5.43 | 0.22 | 0 | 0 | 0 | 0 | 0 | - |
| 40 | T296$^{ECL2}$-L5.43 | 0.21 | 0 | 0 | 0 | 0 | 0 | - |
| 42 | R2.39-S6.36 | 0.21 | 0 | 0 | 0 | 0 | 0 | - |
| 47 | M3.32-P4.59 | 0.20 | 0 | 0 | 0 | 0 | 0 | - |
| 48 | I3.36-S7.42 | 0.20 | 0 | 0 | 0 | 0 | 0 | Contact Definition |
| 50 | R2.39-S2.49 | 0.20 | 0 | 0 | 0 | 0 | 0 | - |
| 55 | L3.48-P6.42 | 0.20 | 0 | 0 | 0 | 0 | 0 | - |
| 58 | L258$^{ICL2}$-K5.68 | 0.20 | 0 | 0 | 0 | 0 | 0 | - |
| 64 | L1.60-Y4.45 | 0.19 | 0 | 0 | 0 | 0 | 0 | - |
| 65 | L3.44-K6.35 | 0.19 | 0 | 0 | 0 | 0 | 0 | - |
| 66 | H3.51-Q5.63 | 0.19 | 0 | 0 | 0 | 0 | 0 | - |

**Table S7. Residue contacts that are important for (A) activation and (B) allosteric modulation of class B GPCRs. (A)** The amino acid identities are listed using the glucagon receptor as a model class B GPCR. $CF_{PDBs}$, $CF_{Active}$ and $CF_{Inactive}$ are the contact frequencies in all PDB, active and inactive structures of class B GPCRs. $\Delta CF_{Activate}$ is the difference in contact frequencies between the active and inactive GPCR structures. **(B)** The amino acid identities are taken from the GLP-1 receptor (PAM) and glucagon receptor (NAM). $CF_{PAM}$, $CF'_{Active}$, $CF_{NAM}$ and $CF'_{Inactive}$ are the contact frequencies in class B GPCR structures bound by the agonist and PAM (PDB: 6VCB), only agonist (PDB: 6B3J), antagonist and NAM (PDB: 5EE7), and only antagonist (PDB: 5XEZ). $\Delta CF_{PAM}$ is the difference in contact frequencies between agonist-PAM and agonist bound GPCR structures, and $\Delta CF_{NAM}$ is the difference in contact frequencies between antagonist-NAM and antagonist bound GPCR structures.

| *Rank* | *Residues* | $CF_{PDBs}$ | $CF_{Active}$ | $CF_{Inactive}$ | $\Delta CF_{Activate}$ | Type |
|---|---|---|---|---|---|---|
| 1 | L3.43-L6.43 | 0.52 | 1.00 | 0 | 1.00 | *Switching* |
| 2 | E3.46-L6.43 | 0.52 | 1.00 | 0 | 1.00 | *Switching* |
| 3 | E3.46-L6.44 | 0.52 | 1.00 | 0 | 1.00 | *Switching* |
| 4 | G3.47-L6.44 | 0.52 | 1.00 | 0 | 1.00 | *Switching* |
| 5 | N5.54-L6.44 | 0.52 | 1.00 | 0 | 1.00 | *Switching* |
| 6 | F5.58-L6.44 | 0.52 | 1.00 | 0 | 1.00 | *Switching* |
| 7 | L3.50-L6.33 | 0.45 | 0 | 1.00 | -1.00 | *Switching* |
| 8 | L3.50-A6.34 | 0.45 | 0 | 1.00 | -1.00 | *Switching* |
| 9 | L3.50-T6.37 | 0.45 | 0 | 1.00 | -1.00 | *Switching* |
| 10 | L3.54-K6.30 | 0.45 | 0 | 1.00 | -1.00 | *Switching* |
| 11 | I5.61-A6.34 | 0.45 | 0 | 1.00 | -1.00 | *Switching* |
| 12 | L6.40-L7.48 | 0.48 | 0 | 1.00 | -1.00 | *Switching* |
| 13 | L6.44-Q7.45 | 0.45 | 0 | 1.00 | -1.00 | *Switching* |
| 14 | I5.57-L6.44 | 0.48 | 0.94 | 0 | 0.94 | *Switching* |
| 15 | L5.69-L6.33 | 0.48 | 0.94 | 0 | 0.94 | *Switching* |
| 16 | P6.42-Q7.45 | 0.48 | 0.94 | 0 | 0.94 | *Switching* |
| 17 | G3.47-T6.37 | 0.42 | 0 | 0.93 | -0.93 | *Switching* |
| 18 | F5.58-L6.38 | 0.45 | 0 | 0.93 | -0.93 | *Switching* |
| 19 | F5.55-V6.46 | 0.48 | 0.88 | 0 | 0.88 | *Switching* |
| 20 | F5.58-L6.40 | 0.45 | 0.88 | 0 | 0.88 | *Switching* |
| 21 | F5.58-V6.46 | 0.45 | 0.88 | 0 | 0.88 | *Switching* |
| 22 | V5.62-T6.37 | 0.45 | 0.88 | 0 | 0.88 | *Switching* |
| 23 | V5.62-L6.40 | 0.45 | 0.88 | 0 | 0.88 | *Switching* |
| 24 | L5.65-L6.33 | 0.45 | 0.88 | 0 | 0.88 | *Switching* |
| 25 | P6.42-L7.48 | 0.45 | 0.88 | 0 | 0.88 | *Switching* |
| 26 | L5.65-A6.34 | 0.55 | 0.12 | 1.00 | -0.88 | Repacking |
| 27 | L3.43-L6.40 | 0.39 | 0 | 0.86 | -0.86 | *Switching* |
| 28 | L3.43-I6.41 | 0.39 | 0 | 0.86 | -0.86 | *Switching* |
| 29 | E3.46-T6.37 | 0.39 | 0 | 0.86 | -0.86 | *Switching* |
| 30 | E3.46-L6.40 | 0.39 | 0 | 0.86 | -0.86 | *Switching* |

**(A)**

| Rank | Residues | $CF_{PAM}$ | $CF'_{Active}$ | $\Delta CF_{PAM}$ | Modulator |
|---|---|---|---|---|---|
| 1 | Y3.40-E6.48 | 0 | 1.00 | -1.00 | PAM |
| 2 | G3.47-L6.43 | 1.00 | 0 | 1.00 | PAM |
| 3 | L3.50-L6.43 | 1.00 | 0 | 1.00 | PAM |
| 4 | L3.50-L6.44 | 1.00 | 0 | 1.00 | PAM |
| 5 | I5.51-V6.46 | 1.00 | 0 | 1.00 | PAM |
| 6 | L5.52-A6.52 | 1.00 | 0 | 1.00 | PAM |
| 7 | N5.54-V6.46 | 1.00 | 0 | 1.00 | PAM |
| 8 | P5.55-L6.44 | 1.00 | 0 | 1.00 | PAM |
| 9 | H6.47-T7.37 | 1.00 | 0 | 1.00 | PAM |
| 10 | H6.47-T7.42 | 1.00 | 0 | 1.00 | PAM |
| 11 | E6.48-E7.38 | 1.00 | 0 | 1.00 | PAM |
| 12 | E6.48-F7.41 | 0 | 1.00 | -1.00 | PAM |
| 13 | I6.50-E7.38 | 1.00 | 0.00 | 1.00 | PAM |
| **Rank** | **Residues** | $CF_{NAM}$ | $CF'_{Inactive}$ | $\Delta CF_{NAM}$ | **Modulator** |
| 1 | L3.43-T6.37 | 1.00 | 0 | 1.00 | NAM |
| 2 | E3.46-I6.41 | 1.00 | 0 | 1.00 | NAM |
| 3 | Y3.49-T6.37 | 0 | 1.00 | -1.00 | NAM |
| 4 | L3.50-K6.30 | 1.00 | 0 | 1.00 | NAM |
| 5 | W5.40-A6.52 | 1.00 | 0 | 1.00 | NAM |
| 6 | R5.44-E6.48 | 1.00 | 0 | 1.00 | NAM |
| 7 | F5.48-A6.52 | 1.00 | 0 | 1.00 | NAM |
| 8 | I5.51-L6.44 | 0 | 1.00 | -1.00 | NAM |
| 9 | H6.47-L7.41 | 0 | 1.00 | -1.00 | NAM |
| 10 | H6.47-Q7.45 | 1.00 | 0 | 1.00 | NAM |
| 11 | V6.50-K7.34 | 1.00 | 0 | 1.00 | NAM |
| 12 | V6.50-D7.38 | 1.00 | 0 | 1.00 | NAM |

**(B)**

**Table S8. Top ranked exception residue contacts predicted in the Potts model that are important for activation and allosteric modulation of (A) class A (using residue index in β1-adrenoreceptor) and (B) class B (using residue index in glucagon receptor) GPCRs**. Only residue pairs with at least five residues apart in the numbering scheme were considered. $d_{PDBs}$, $d_{Active}$ and $d_{Inactive}$ are the Cβ-Cβ distances in all PDB, active and inactive class A GPCR structures. $d_{PAM}$ and $d_{NAM}$ are the Cβ-Cβ distances in agonist-PAM and antagonist-NAM bound GPCR structures.

| Rank | Residues | $d_{PDBs}$ | $d_{Active}$ | $d_{Inactive}$ | $d_{PAM}$ | $d_{NAM}$ |
|---|---|---|---|---|---|---|
| 4 | Y3.51-S3.56 | 8.88 ± 0.76 | 8.68 ± 0.64 | 8.97 ± 0.80 | 9.07 ± 0.57 | 10.05 ± 0.82 |
| 11 | A2.49-W4.50 | 10.12 ± 2.20 | 10.69 ± 2.53 | 9.85 ± 1.98 | 8.16 ± 0.20 | 8.84 ± 0.29 |
| 14 | C3.44-V5.57 | 9.41 ± 1.91 | 9.61 ± 1.82 | 9.31 ± 1.94 | 9.02 ± 0.88 | 9.16 ± 2.76 |
| 18 | W4.50-P7.50 | 21.15 ± 2.23 | 20.43 ± 2.49 | 21.56 ± 1.96 | 19.67 ± 0.35 | 20.73 ± 0.07 |
| 20 | $C198^{ECL2}$-N5.35 | 17.51 ± 3.64 | 18.79 ± 3.54 | 17.00 ± 3.45 | 17.98 ± 0.24 | 17.49 ± 0.34 |
| 28 | Y3.51-F5.56 | 8.96 ± 2.49 | 7.53 ± 2.74 | 9.66 ± 2.01 | 9.63 ± 0.36 | 9.68 ± 0.58 |
| 29 | T3.42-W4.50 | 9.60 ± 2.19 | 10.18 ± 2.37 | 9.27 ± 2.05 | 8.51 ± 0.66 | 9.18 ± 0.08 |
| 30 | D3.49-$Y149^{ICL2}$ | 10.30 ± 2.78 | 10.14 ± 2.71 | 10.35 ± 2.75 | 8.50 ± 0.24 | 8.27 ± 0.91 |
| 34 | S3.30-W6.48 | 18.68 ± 2.82 | 20.80 ± 2.91 | 17.54 ± 2.02 | 18.96 ± 0.19 | 17.62 ± 0.27 |
| 37 | R3.50-N7.45 | 19.98 ± 1.27 | 18.81 ± 0.83 | 20.59 ± 1.02 | 18.40 ± 0.51 | 20.38 ± 0.24 |
| 39 | $C198^{ECL2}$-$T203^{ECL2}$ | 15.31 ± 3.16 | 16.70 ± 3.41 | 14.69 ± 2.76 | 14.69 ± 0.53 | 14.61 ± 0.09 |
| 41 | D3.32-Y7.43 | 10.04 ± 2.42 | 9.80 ± 1.63 | 10.17 ± 2.69 | 9.25 ± 0.26 | 9.04 ± 0.18 |
| 44 | Y5.58-P7.50 | 21.93 ± 2.74 | 20.63 ± 2.75 | 22.62 ± 2.51 | 19.37 ± 0.23 | 22.78 ± 3.84 |
| 48 | F2.42-$M153^{ICL2}$ | 11.83 ± 1.88 | 11.64 ± 1.91 | 11.91 ± 1.86 | 11.08 ± 0.75 | 11.32 ± 0.80 |
| 53 | N6.61-D7.32 | 13.54 ± 2.91 | 14.40 ± 3.24 | 13.04 ± 2.60 | 15.00 ± 0.20 | 15.10 ± 0.71 |
| 56 | L4.59-Y5.38 | 11.79 ± 2.62 | 11.68 ± 2.56 | 11.91 ± 2.61 | 11.33 ± 0.58 | 10.12 ± 2.73 |
| 57 | I6.40-P6.50 | 17.34 ± 0.79 | 17.14 ± 0.76 | 17.48 ± 0.75 | 17.64 ± 0.37 | 17.66 ± 0.37 |
| 60 | F5.47-L6.49 | 10.16 ± 3.36 | 10.59 ± 4.09 | 9.93 ± 2.95 | 7.68 ± 0.23 | 10.18 ± 2.65 |
| 61 | L2.46-S7.46 | 9.59 ± 0.80 | 9.56 ± 0.81 | 9.63 ± 0.80 | 9.00 ± 0.30 | 10.03 ± 0.62 |
| 62 | S4.53-N7.49 | 18.56 ± 2.70 | 17.82 ± 2.28 | 18.99 ± 2.83 | 15.96 ± 0.38 | 17.23 ± 0.66 |

**(A) Class A**

| Rank | Residues | $d_{PDBs}$ | $d_{Active}$ | $d_{Inactive}$ | $d_{PAM}$ | $d_{NAM}$ |
|---|---|---|---|---|---|---|
| 2 | S2.49-W4.50 | 8.31 ± 0.39 | 8.40 ± 0.49 | 8.20 ± 0.21 | 8.11 | 8.22 |
| 7 | Q3.33-L4.66 | 16.47 ± 0.69 | 16.54 ± 0.69 | 16.41 ± 0.72 | 16.13 | 16.09 |
| 19 | W295$^{ECL2}$-L5.43 | 11.24 ± 1.12 | 11.46 ± 0.91 | 11.05 ± 1.33 | 11.01 | 11.44 |
| 21 | H2.43-L6.40 | 12.80 ± 1.68 | 13.66 ± 0.54 | 11.41 ± 0.82 | 13.34 | 10.94 |
| 23 | L6.44-S7.42 | 11.52 ± 2.29 | 13.54 ± 0.92 | 9.19 ± 0.44 | 12.45 | 8.72 |
| 27 | Q3.33-E6.48 | 15.74 ± 2.12 | 16.99 ± 1.63 | 14.04 ± 1.07 | 16.61 | 13.35 |
| 30 | F5.58-L6.33 | 13.43 ± 1.59 | 13.55 ± 1.83 | 13.24 ± 1.36 | 13.27 | 13.63 |
| 32 | N5.54-L6.43 | 10.74 ± 1.77 | 9.57 ± 0.54 | 11.81 ± 1.69 | 9.48 | 11.39 |
| 33 | A2.54-L7.39 | 13.41 ± 0.85 | 13.59 ± 0.86 | 13.24 ± 0.83 | 12.06 | 12.86 |
| 35 | K2.53-L5.43 | 14.04 ± 1.04 | 14.14 ± 1.08 | 13.92 ± 1.06 | 14.51 | 13.40 |
| 40 | T296$^{ECL2}$-L5.43 | 13.06 ± 1.34 | 13.33 ± 1.41 | 12.76 ± 1.29 | 12.42 | 13.10 |
| 42 | R2.39-S6.36 | 14.53 ± 3.29 | 16.94 ± 1.27 | 11.28 ± 0.94 | 16.89 | 10.25 |
| 47 | M3.32-P4.59 | 16.38 ± 0.42 | 16.24 ± 0.52 | 16.57 ± 0.18 | 16.08 | 16.53 |
| 48 | I3.36-S7.42 | 10.04 ± 1.18 | 10.41 ± 1.33 | 9.73 ± 0.84 | 9.07 | 10.15 |
| 50 | R2.39-S2.49 | 16.13 ± 0.54 | 15.83 ± 0.29 | 16.52 ± 0.50 | 15.81 | 15.82 |
| 55 | L3.48-P6.42 | 17.02 ± 1.32 | 17.06 ± 0.45 | 17.06 ± 1.91 | 16.37 | 16.14 |
| 58 | L258$^{ICL2}$-K5.68 | 18.84 ± 2.32 | 19.51 ± 2.93 | 18.15 ± 1.17 | 19.74 | 17.92 |
| 64 | L1.60-Y4.45 | 23.56 ± 7.80 | 22.25 ± 0.75 | 25.16 ± 11.61 | 22.20 | 21.07 |
| 65 | L3.44-K6.35 | 21.44 ± 2.62 | 23.80 ± 0.72 | 18.68 ± 0.40 | 23.40 | 18.22 |
| 66 | H3.51-Q5.63 | 11.16 ± 0.55 | 11.45 ± 0.51 | 10.85 ± 0.41 | 11.01 | 10.07 |

**(B) Class B**

**Table S9. Comparison of residue contacts that are important for activation of class A GPCRs between Cvicek et al., 2016 (15), Zhou et al., 2019 (16) and our work. (A)** 10 residue contacts from our lists (**Figure 2**) that also appeared in the Native activation "hot-spot" residues (NACHOs) (15). **(B)** 9 residue contacts from our lists (**Figure 2**) that also appeared in the common activation pathway of class A GPCRs (16). 5 previously identified switching contacts that became repacking contacts are marked with *. **(C)** 20 additional residue contacts that were found important in the activation of class A GPCRs. The repacking residue contacts are shown in regular fonts, while the switching residue contacts are bold.

| **A** (10) | **B** (9) | **C** (20) | |
|---|---|---|---|
| 1.53-7.53 | 1.53-7.53* | 1.53-7.49 | 3.39-7.49 |
| 3.43-6.41 | 3.43-6.41* | 2.53-3.39 | 5.54-6.40 |
| **3.46-6.37** | **3.46-6.37** | 2.46-7.53 | 5.54-6.44 |
| **3.50-6.34** | 3.50-6.37* | 2.47-7.49 | 5.61-6.33 |
| 3.43-7.49 | 3.43-7.49* | 2.50-7.50 | 5.61-6.34 |
| **3.46-7.53** | **3.46-7.53** | **3.39-6.44** | 5.61-6.37 |
| **5.58-6.40** | **5.58-6.40** | 3.47-6.37 | 5.65-6.31 |
| **5.62-6.37** | **5.62-6.37** | 3.47-6.41 | **5.65-6.33** |
| 6.36-7.53 | 6.40-7.49* | **3.50-6.34** | 6.36-7.52 |
| 6.40-7.49 | | 3.39-7.46 | 6.46-7.41 |

**References**


1. S. R. Eddy, Accelerated Profile HMM Searches. *PLoS Computational Biology* **7**, e1002195 (2011).
2. A. Haldane, Flynn, W.F., He, P., Vijayan, R.S., Levy, R.M., Structural propensities of kinase family proteins from a Potts model of residue covariation. *Protein Sciences* **25**, 1378-1384 (2016).
3. A. Haldane, Flynn, W.F., He, P., Levy, R.M., Coevolutionary Landscape of Kinase Family Proteins: Sequence Probabilities and Functional Motifs. *Biophysical Journal* **114**, 21-31 (2018).
4. F. Morcos, Pagnani, A., Lunt, B., Bertolino, A., Marks, D.S., Sander, C., Zecchina, R., Onuchic, J.N., Hwa, T., Weigt, M., Direct-coupling analysis of residue coevolution captures native contacts across many protein families. *Proc Natl Acad Sci USA* **108**, E1293-1301 (2011).
5. R. M. Levy, Haldane, A., Flynn, W.F., Potts Hamiltonian models of protein covariation, free energy landscapes, and evolutionary fitness. *Current Opinion in Structural Biology* **43**, 55-62 (2017).
6. M. Weigt, White, R.A., Szurmant, H., Hoch, J.A., Hwa, T., Identification of direct residue contacts in protein-protein interaction by message passing. *Proc Natl Acad Sci USA* **106**, 67-72 (2008).
7. A. Haldane, Levy, R.M., Mi3-GPU: MCMC-based Inversing Ising Inference on GPUs for protein covariation analysis. *Computer Physics Communications* **107312** (2020).
8. M. Ekeberg, C. Lövkvist, Y. Lan, M. Weigt, E. Aurell, Improved contact prediction in proteins: Using pseudolikelihoods to infer Potts models. *Physcial Review E* **87**, 012707 (2013).
9. H. M. Berman, Westbrook, J., Feng, Z., Gilliland, G., Bhat, T.N., Weissig, H., Shindyalov, I.N., Bourne, P.E., The Protein Data Bank. *Nucleic Acids Research* **28**, 235-242 (2000).
10. S. K. Burley, Berman, H.M., Bhikadiya, C., Bi, C., Chen, L., Di Costanzo, L., Christie, C., Dalenberg, K., Duarte, J.M., Dutta, S., Feng, Z., Ghosh, S., Goodsell, D.S., Green, R.K., Guranovic, V., Guzenko, D., Hudson, B.P., Kalro, T., Liang, Y., Lowe, R., Namkoong, H., Peisach, E., Periskova, I., Prlic, A., Randle, C., Rose, A., Rose, P., Sala, R., Sekharan, M., Shao, C., Tan, L., Tao, Y.P., Valasatava, Y., Voigt, M., Westbrook, J., Woo, J., Yang, H., Young, J., Zhuravleva, M., Zardecki, C., RCSB Protein Data Bank: biological macromolecular structures enabling research and education in fundamental biology, biomedicine, biotechnology and energy. *Nucleic Acids Research* **47**, D464-D474 (2019).
11. G. Pándy-Szekeres, Munk, C., Tsonkov, T.M., Mordalski, S., Harpsøe, K., Hauser, A.S., Bojarski, A.J., Gloriam, D.E., GPCRdb in 2018: adding GPCR structure models and ligands. *Nucleic Acids Research* **46**, D440-D446 (2018).
12. V. Isberg, de Graaf, C., Bortolato, A., Cherezov, V., Katritch, V., Marshall, F.H., Mordalski, S., Pin, J.P., Stevens, R.C., Vriend, G., Gloriam, D.E., Generic GPCR Residue Numbers- Aligning Topology Maps While Minding The Gaps. *Trends in Pharmacological Sciences* **36**, 22-31 (2015).
13. R. van der Kant, Vriend, G., Alpha-bulges in G protein-coupled receptors. *International Journal of Molecular Sciences* **15**, 7841-7864 (2014).
14. Q. Zhou *et al.*, Common activation mechanism of class A GPCRs. *Elife* **19**, e50279 (2019).

15. V. Cvicek, Goddard, W.I., Abrol, R., Structure-Based Sequence Alignment of the Transmembrane Domains of All Human GPCRs: Phylogenetic, Structural, and Functional Implications. *PLoS Computational Biology* **12**, e1004805 (2016).
16. Q. Zhou, Yang, D., Wu, M., Guo, Y., Guo, W., Zhong, L., Cai, X., Dai, A., Jang, W., Shakhnovich, E.I., Liu, Z.J., Stevens, R.C., Lambert, N.A., Babu, M.M., Wang, M.W., Zhao, S., Common activation mechanism of class A GPCRs. *Elife* **19**, e50279 (2019).
